# miR319-targeted *TCP4/LANCEOLATE* directly regulates *OVATE* and auxin responses to modulate tomato gynoecium patterning and fruit morphology

**DOI:** 10.1101/2022.03.15.484181

**Authors:** Airton Carvalho, Mateus H. Vicente, Leticia F. Ferigolo, Eder M. Silva, Matan Levy, Lazaro E.P. Peres, Robert Sablowski, Carla Schommer, Naomi Ori, Fabio T.S. Nogueira

**Author notes:** These authors contributed equally for this work (Airton Carvalho, Mateus H. Vicente, Leticia F. Ferigolo). To whom correspondence should be addressed: Fabio T.S. Nogueira.

## Abstract

Diversity in fruit morphology is one of the hallmarks of varietal differences among modern cultivars of fruit-bearing crops. As evolutionarily related organs, fruits and leaves share developmental processes, but there are surprisingly few connections between regulatory pathways for fruit and leaf development. Here, we show the regulation of the leaf development-associated TEOSINTE BRANCHED1/CYCLOIDEA/PCF (TCP) TCP4/LANCEOLATE (TCP4/LA) transcription factor by the microRNA319 (miR319) is crucial for gynoecium patterning and establishment of fruit morphology. Loss of miR319 regulation leads to a premature, ectopic *TCP4/LA* expression during gynoecium patterning, which results in elongated fruits, resembling *ovate* mutants. *TCP4/LA* modulates tomato fruit development and morphology partially by directly repressing *OVATE* expression as early as 5-8 days post-inflorescence (dpi) flower buds. Furthermore, miR319-targeted *CINCINNATA-like TCP4/LANCEOLATE* controls auxin responses in developing flower buds by directly binding to the *SlYUCCA4* promoter. Modulation of auxin biosynthesis by *TCP4/LA* is shared with other *CINCINNATA-like TCPs* during *Arabidopsis* gynoecium patterning. Our study defines a novel miRNA-based molecular link between *OVATE*, a fundamental gene associated with tomato domestication, and auxin responses in the control of fruit development and morphology. Given the striking variation in fruit shape among members of the Solanaceae family, fine-tuning regulation of gene expression by miRNA coupled with modulation of hormone dynamics may be a common driver in the evolution of fruit-shape diversity.

## INTRODUCTION

Diversity in fruit morphology is one of the hallmarks of varietal differences among modern cultivars of tomatoes, peppers and other fruit-bearing crops. Fruit morphogenesis is determined by the coordination of cell division and expansion, which are fundamental processes required for the development of all plant organs (van der Knaap *et al*., 2014; Snouffer *et al*., 2020). Therefore, unravelling the molecular basis of fruit morphogenesis not only provide insights into general mechanisms of plant organogenesis, but also help to understand the process of domestication, and perhaps facilitate breeding new cultivars for crop improvement.

Fruit shape diversity is a result of the activities of key regulatory genes along with different patterns of anisotropic growth and hormone responses (Seymour *et al*., 2013; Eldridge *et al*., 2016). In tomatoes and melons, for instance, the diversity in fruit morphology is a result of thousands of years of selection under cultivation, which resulted in the accumulation of specific mutations. Quantitative trait loci (QTL) studies have revealed a number of loci that control fruit morphology in tomato, including *fw2.2, fw3.2, ovate, sun, locule-number* (*lc*), and *fasciated* (*fas*)(van der Knaap *et al*., 2014). For instance, the loss of *OVATE* function, the founding member of the conserved OVATE Family Protein (OFP), leads to elongated fruit phenotype in tomato (Liu *et al*., 2002; Rodriguez *et al*., 2011). However, the morphological fruit complexity exhibited by several crops suggests that additional genetic factors exist and are fundamental for coordinating cell division and expansion, patterning of the gynoecium, and specification of organ shape and seed dispersal. Therefore, new strategies in addition to classic QTL studies are necessary to further dissect genetic networks shared by all higher plants to control fruit morphogenesis (van der Knaap & Østergaard, 2018).

Only few genes controlling fruit shape and development have been directly associated with leaf development (Wu *et al*., 2018). This is surprising, given that carpels and leaves are evolutionarily related organs, as carpels are thought to be modified leaves. Thus, developmental pathways and genes that play crucial roles in patterning both organs are likely conserved (Scutt *et al*., 2006; van der Knaap *et al*., 2014). In fact, this idea has been supported by early studies from *Arabidopsis* such as the transformation of floral organs into carpelloid leaf-like organs in a triple mutant lacking the ABC homeotic functions, the transformation of vegetative leaves into floral organs by the ectopic expression of *SEPALLATA* and other floral homeotic genes, and the increase in replum size by the misexpression of *BREVIPEDICELLUS* (a leaf development associated gene) in the ovary (Bowman *et al*., 1991; Honma & Goto, 2001; Pelaz *et al*., 2001; Østergaard, 2009).

Some of the genes with common functions during leaf and gynoecium/fruit development are transcription factors (TFs), which control the timely progression of organ patterning and development through their interaction with hormone dynamics. Phytohormones such as auxin and gibberellin are intricately involved in the determination of gynoecium morphology and functionality (Alonso-Cantabrana *et al*., 2007; Fuentes *et al*., 2012; Moubayidin & Østergaard, 2014). Post-transcriptional regulation of TFs mediated by microRNAs (miRNAs) is yet another fundamental layer of controlling proper leaf and fruit development (Juarez *et al*., 2004; Ori *et al*., 2007; Xing *et al*., 2013; Silva *et al*., 2014). However, the relevance of miRNA-dependent mechanisms associated with leaf development and their interplay with phytohormones in gynoecium patterning and fruit development is still poorly understood. In both tomato and *Arabidopsis*, the miR319-mediated repression of a sub-group of the <u>T</u>EOSINTE BRANCHED1/<u>C</u>YCLOIDEA/<u>P</u>CF (TCP) family of TFs (called *CINCINNATA-like TCPs* or *CIN-TCPs*) is crucial to proper leaf development through the modulation of the balance between cell proliferation and differentiation (Palatnik *et al*., 2003; Ori *et al*., 2007). The loss of miR319 regulation of the *CIN-TCP LANCEOLATE* (*LA*) leads to simpler leaves in the semi-dominant tomato *Lanceolate* (*La*) mutant (Ori *et al*., 2007). In *Arabidopsis*, miR319-based regulation of *TCP4* (homologue of *LANCEOLATE*) is critical for petal and stamen development (Nag *et al*., 2009), and the overexpression of miR319a in the dominant *Jaw-D* mutant leads to leaf and silique developmental defects (Palatnik *et al*., 2003). Nevertheless, the underlying mechanisms by which miR319-targeted *CIN-TCPs* modulate fruit morphology and their relevance to seed production remain unknown.

One possibility is that *TCPs* modulate fruit morphology through the hormone auxin. Auxin controls cell proliferation and maturation in a developmental context-dependent manner. In tomato, auxin treatment before anthesis led to elongated ovaries and modified fruit shape (Wang *et al*., 2019). Arabidopsis plants expressing a dominant, repressive form of the *TCP15* display up-regulation of the auxin biosynthesis genes *YUCCA1* and *YUCCA4*, suggesting that *TCP15* is a negative regulator of auxin biosynthesis (Lucero *et al*., 2015). Similarly, *TCP3* inhibits auxin response by inducing IAA3 (INDOLE-3-ACETIC ACID INDUCIBLE3)/ SHY2 (SHORT HYPOCOTYL2), a negative regulator of auxin signalling (Koyama *et al*., 2010). By contrast, *TCP4* induces auxin response by activating *YUCCA5* in the hypocotyl (Challa et al., 2016). Even though the regulation of auxin response by TCP proteins has been investigated, how miR319-targeted *CIN-TCPs* connect auxin homeostasis and fruit morphology is poorly understood.

Here we show that miR319-dependent regulation of *TCP4*/*LANCEOLATE* is fundamental to modulate tomato fruit development. Semi-dominant *Lanceolate* mutant showed premature carpel differentiation, resulting in elongated fruits resembling classical *ovate* fruit phenotypes, but with a more constrict fruit phenotype. The genetic and molecular results shown in this study are consistent with LA directly repressing the activity of the major domestication fruit shape gene *OVATE* during early carpel patterning. Our data also indicate that miR319-targeted *LA* controls auxin homeostasis during early gynoecium patterning by directly repressing *YUCCA4*, thus impacting final fruit shape and seed production. This work defines a leaf development-associated miRNA-controlled module that integrates developmental and hormone signalling pathways in the control of fruit development and morphology.

## RESULTS

### Modulation of *TCP4*/*LANCEOLATE* expression is crucial to control gynoecium and fruit shape

*Arabidopsis MIR319* loss-of-function flowers present narrow petals as well as undeveloped anthers (Nag *et al*., 2009), suggesting that the miR319/*CIN-TCP* regulatory hub is important not only in leaf development (Ori *et al*., 2007; Ben-Gera & Ori, 2012; Burko *et al*., 2013), but also in some aspects of the flower development. To test whether the miR319 regulatory hub contributes to fruit growth, we initially evaluated fruit phenotype of tomato mutant plants from an indeterminate cultivar (accession LA0335) that displays simpler leaves, and is catalogued as a *Lanceolate* (*La*) allele in the TGRC (http://tgrc.ucdavis.edu). The classical semi-dominant *La* mutation causes the dose-dependent and gradual conversion of compound tomato leaves into small simple ones, as well smaller sepals and fruits (Stettler, 1964). Previously, we have cloned the *Lanceolate* (*La-1*) allele from LA0335 plants, which is characterized by a synonymous mutation in the miR319 binding site that results in *LA* de-repression (Silva *et al*., 2019). Because homozygous *La* mutants are rarely viable (Stettler, 1964; Silva *et al*., 2019), we characterized the semi-dominant *La-1/+* (LA0335) flowers and fruits (Figure 1a, b). *La-1/+* (LA0335) flowers displayed shorter sepals when compared with non-*La-1/+*wild-type (WT) LA0335 segregant (Figure 1b), similarly to Arabidopsis *miR319* mutants (Nag *et al*., 2009). Importantly, *La-1/+* (LA0335) fruits are elongated and show absent or malformed septum and a rudimental placenta (Figure 1a), which indicate developmental defects during carpel development. In fact, all pre-anthesis *La-1/+* (LA0335) flowers presented carpels with a medial constriction in the ovary (Figure 1b, arrowhead). Supporting these observations, *La-1/+* (LA0335) ovaries and fruits showed significant changes in shape (Figure S1a).

**Figure 1.**
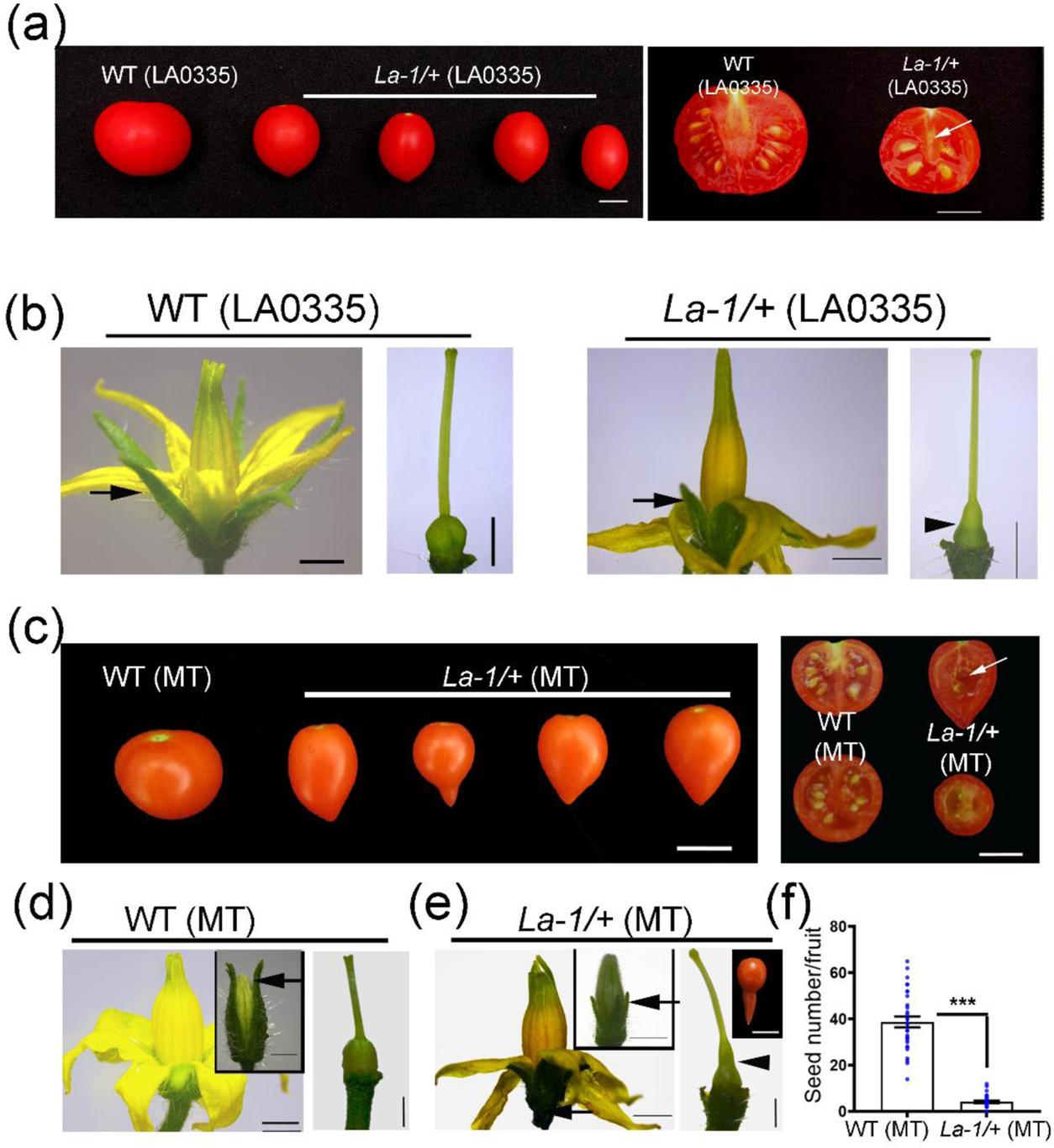
*LANCEOLATE* de-repression leads to reproductive defects in tomato. (a) Left panel: representative elongated and small fruit phenotypes of *La-1/+* (LA0335) fruits. Right panel: modified inner architecture of *La-1/+* (LA0335) fruits indicated by absent or malformed septum and a rudimental placenta. (b) Normal sepals and carpel of WT (LA0335) flowers compared with the smaller sepals of *La-1/+* (LA0335) flowers (arrow) and their altered carpel morphology indicated by the medial constriction (arrowhead). (c) Left panel: representative fruits phenotypes of WT (MT) and *La-1/+* (MT) plants. Right panel: modified inner architecture of *La-1/+* (MT) fruits indicated by absent or malformed septum and a rudimental placenta. (d) WT (MT) flower and carpels. (e) Shorter sepals and constricted ovaries (arrow) of *La-1/+* (MT) similar to *La-1/+* (LA0335) flowers. Extreme *La-1/+* (MT) fruit phenotype showing the medial constriction observed in the mutant carpels (inset). (f) Seed production per fruit of *La-1/+* and WT (MT) (\*\*\**P*<0.001, according to Student’s t-test two-tailed). Values are means ± s.e. Scale bars: 1 cm (a, c), 2 mm (b, d, e).

To determine whether carpel and fruit phenotypes of *La-1/+* (LA0335) plants were reproducible in distinct tomato cultivars, we out-crossed *La-1/+* (LA0335) with tomato cv. Micro-Tom (MT) and M82 plants. The wild-type F1 progeny from the crosses produced normal carpel and fruits, but the F1 offspring with the *La-1* mutation (Silva *et al*., 2019) displayed modified ovaries and elongated fruits (Figure S2a-d), similarly to *La-1/+* (LA0335) plants. These observations indicate that the miR319-mediated *LA* regulation is required for proper carpel and fruit development in the hybrid backgrounds as well.

We have previously introgressed the *La-1* allele from LA0335 into tomato cv. MT (Silva *et al*., 2019), and the *La-1/+* (MT) plants also exhibited elongated fruits with malformed septum and also a rudimental placenta (Figure 1c). Interestingly, several *La-1/+* (MT) fruits displayed a more extreme fruit phenotype (Figure 1e, inset), exhibiting a medial constriction associated with an increase in fruit length (∼20.0 %) and reduced locule number (Figure S1d). Similar to *La-1/+* (LA0335) (Figure 1a, b), flowers of *La-1/+* (MT) display short sepals and narrowing growth of the medial region of the ovaries (Figure 1e). The significant shape changes observed in ovaries and fruits of *La-1/+* (MT) (Figure S1b-d) likely led to lower seed production when compared to wild-type MT fruits (Figure 1f). However, it is also possible that low seed production might impact final fruit development. Taken together, these observations indicate that miR319-based regulation of *LANCEOLATE* is required for proper fruit development and seed production in different tomato cultivars.

Because of the extreme fruit phenotypes observed in *La-1/+* (MT) plants (Figure 1e, inset), we characterized in greater detail the fruit morphology of these plants. We measured the number of cell layers in the wall of ovaries at 0 days post-anthesis (DPA) and in the pericarp of 16-DPA fruits. Cell divisions promoting cell layer formation in the fruit pericarp ceases at 16 DPA in most tomato cultivars (Azzi et al., 2015). Interestingly, 0-DAP ovary wall from *La-1/+* (MT) plants was thicker, showing more cell layers than MT plants (Figure 2a, b). This observation suggests a precocious growth of the ovary wall in *La-1/+* (MT) ovaries. On the other hand, 16-DPA *La-1/+* (MT) fruits showed a thinner pericarp, with a reduced number of cell layers (Figure 2c, d), which indicates that cell divisions after fruit set were impaired in *La-1/+* (MT) fruits.

**Figure 2.**
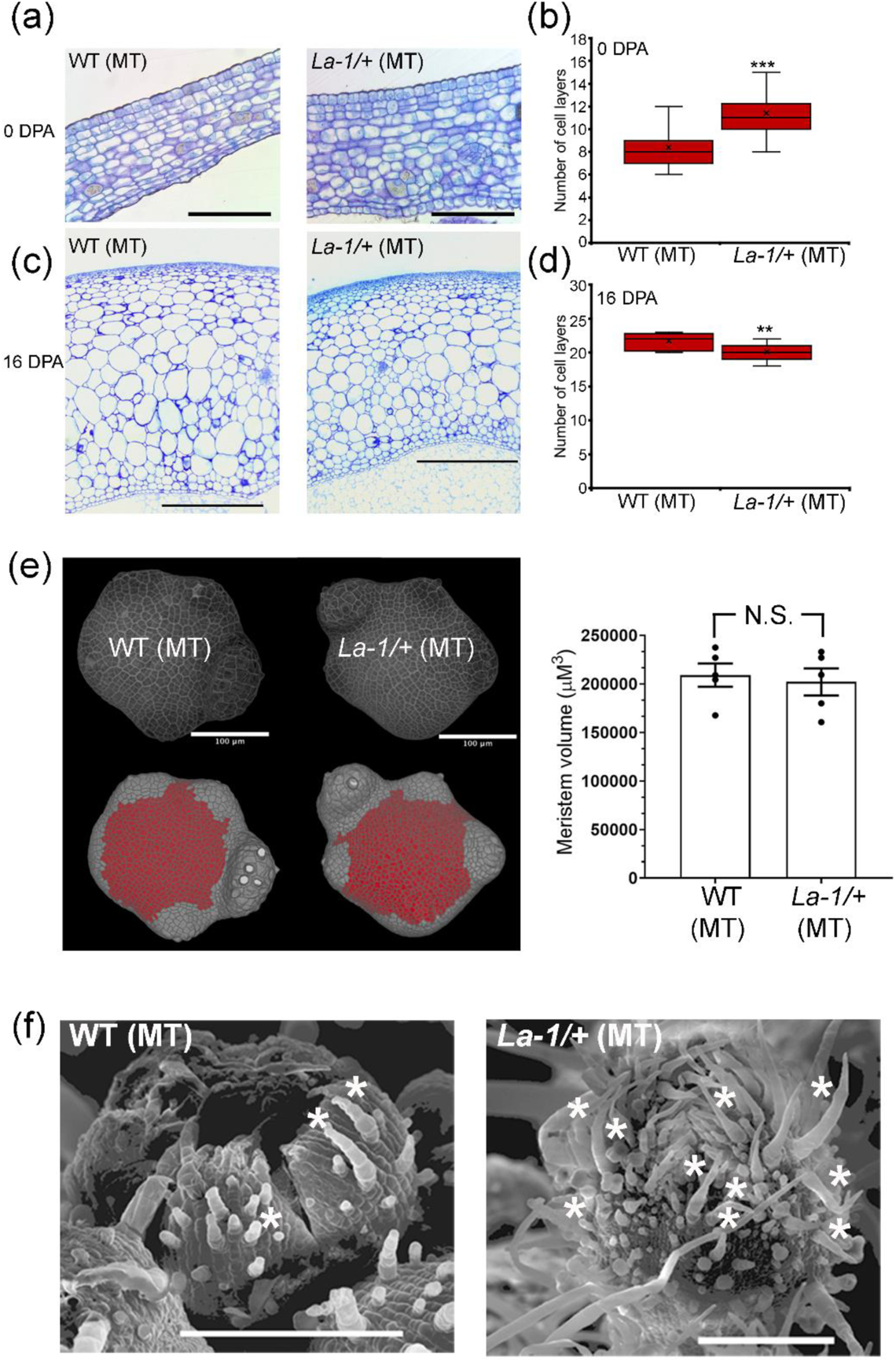
**The loss of miR319 regulation by *TCP4/LA* leads to premature gynoecium patterning.** (a-d) Ovary wall of *La-1/+* (MT) carpels at 0 days post-anthesis (DPA), and fruit pericarp at 16 DPA. At 16 DPA (c), the *La-1/+* (MT) fruit pericarp becomes thinner than WT (MT). (b, d) Cell layers counting of ovary walls showed in (a) at 0 DPA (\*\*\**P*<0.001) and (c) at 16 DPA (\*\**P*<0.01). The black line in the red box indicates the median, the red box indicates the quartiles and the whiskers indicate the minimum and maximum layers. Scale bars: 100 µm (a), 1 mm (b). (e) Right panel: 3D reconstructions from confocal stacks of floral apices of WT (MT) and *La-1/+* (MT) plants stained by the modified pseudo-Schiff propidium iodide (mPS-PI) method (Bencivenga et al., 2016; Serrano-Mislata et al., 2017). Floral meristem domes are shown by their respective epidermal L1 meristematic areas in red. The measured region in red corresponds to the meristem epidermis determined on a 3D image using landmarks placed on the bud boundaries. Asterisks indicate sepal primordia. Left panel: Volume measurements of the meristem epidermis detected no significant differences between genotypes. (f) Scanning electron microscopy of closed buds at 2-5 days post floral initiation (dpi) shows a higher density of trichomes (asterisks) in the *La-1/+* (MT) sepals. Scale bars: 2 mm.

Reduced pericarp growth at 16 DPA of *La-1/+* (MT) fruits may be due the expressive decreasing in seed number of *La-1/+* (MT) fruits (Figure 1f), which could lead to lower auxin biosynthesis and transport from developing seeds into the pericarp (Pattison & Catalá, 2012b; Pattison *et al*., 2015). This is in agreement with more cell divisions observed in the pericarp of parthenocarpic fruits induced by auxin (Serrani *et al*., 2008).

Given that *La* mutants often display vegetative meristem maintenance defects (Ori *et al*., 2007; Silva *et al*., 2019), we hypothesized the modifications in the gynoecium patterning and fruit growth in *La-1/+* plants (Figure 1) might be a result of early alterations in the floral meristem size, as result of a higher rate of cell division or precocious tissue differentiation. To test this hypothesis, we measured the epidermal layer of the floral meristem dome as a proxy for meristem size of WT (MT) and *La-1/+* (MT) inflorescences. A modified Pseudo-Schiff Propidium Iodide (mPS-PI) method coupled with confocal imaging and 3D segmentation of tomato floral meristems (Bencivenga *et al*., 2016; Serrano-Mislata *et al*., 2017) showed no obvious difference between WT (MT) and *La-1/+* (MT) meristem size (Figure 2e). This finding indicates the alterations in carpel patterning of *La-1/+* (MT) flowers occur later in flower development. Trichomes, a marker of leaf and sepal differentiation (Hagemann & Gleissberg, 1996; Matías-Hernández *et al*., 2016), emerged from both WT and *La-1/+* (MT) sepals. However, *La-1/+* (MT) flowers displayed increased trichome density in the sepals at similar developmental stages as WT (MT) (Figure 2f), suggesting premature differentiation of *La-1/+* (MT) floral organs. Our results suggest that floral meristem in *La-1/+* (MT) might differentiate earlier than WT (MT), although more cell layers in *La-1/+* (MT) ovary wall at 0 DPA (Figure 2a-d) indicate more cell divisions during the pre-anthesis phase of gynoecium patterning.

A sufficient period of morphogenetic competence is necessary for leaf complexity, which is reinforced by the miR319-dependent repression of *TCP4*/*LANCEOLATE* (*LA*) (Ori *et al*., 2007; Shleizer-Burko *et al*., 2011a). In developing flower whorls, the expression pattern of *CIN-TCPs* and miR319 is less understood, although the tomato homologue in *Arabidopsis*, *TCP4*, is lowly expressed in the ovary wall (Sarvepalli and Nath, 2011). According to the transcriptome analyses of *S. pimpinellifolium* accession LA1589 (Wang *et al*., 2019), *LA* is lowly expressed in floral buds at 4-6 days post floral initiation (dpi), and increases its expression at later developmental stages of flower development (Figure S3). We then analyzed in more details *LA* and miR319 spatiotemporal expression in developing flowers during pre-anthesis by *in situ* hybridization. We analysed two developmental phases (P): from the emergence of the carpel primordia until ovary closure (5-8 days post-inflorescence, or dpi; termed hereafter developmental phase P1), and carpel growth and maturation, with the presence of ovules (> 10 dpi, P2) (van der Knaap *et al*., 2014). At P1 developing gynoecium, *LA* transcripts were detected at low levels in WT (MT) petal, sepal, and anther primordia, but they were undetectable in the innermost tissues of the gynoecium (Figure 3a). At P2 developing gynoecium, *LA* was expressed in the anthers, developing ovules, placenta margins, and stigma (Figure 3b). Interestingly, *LA* transcripts accumulated only in the outermost cell layer of the ovule integument (Figure 3c, arrowheads).

**Figure 3.**
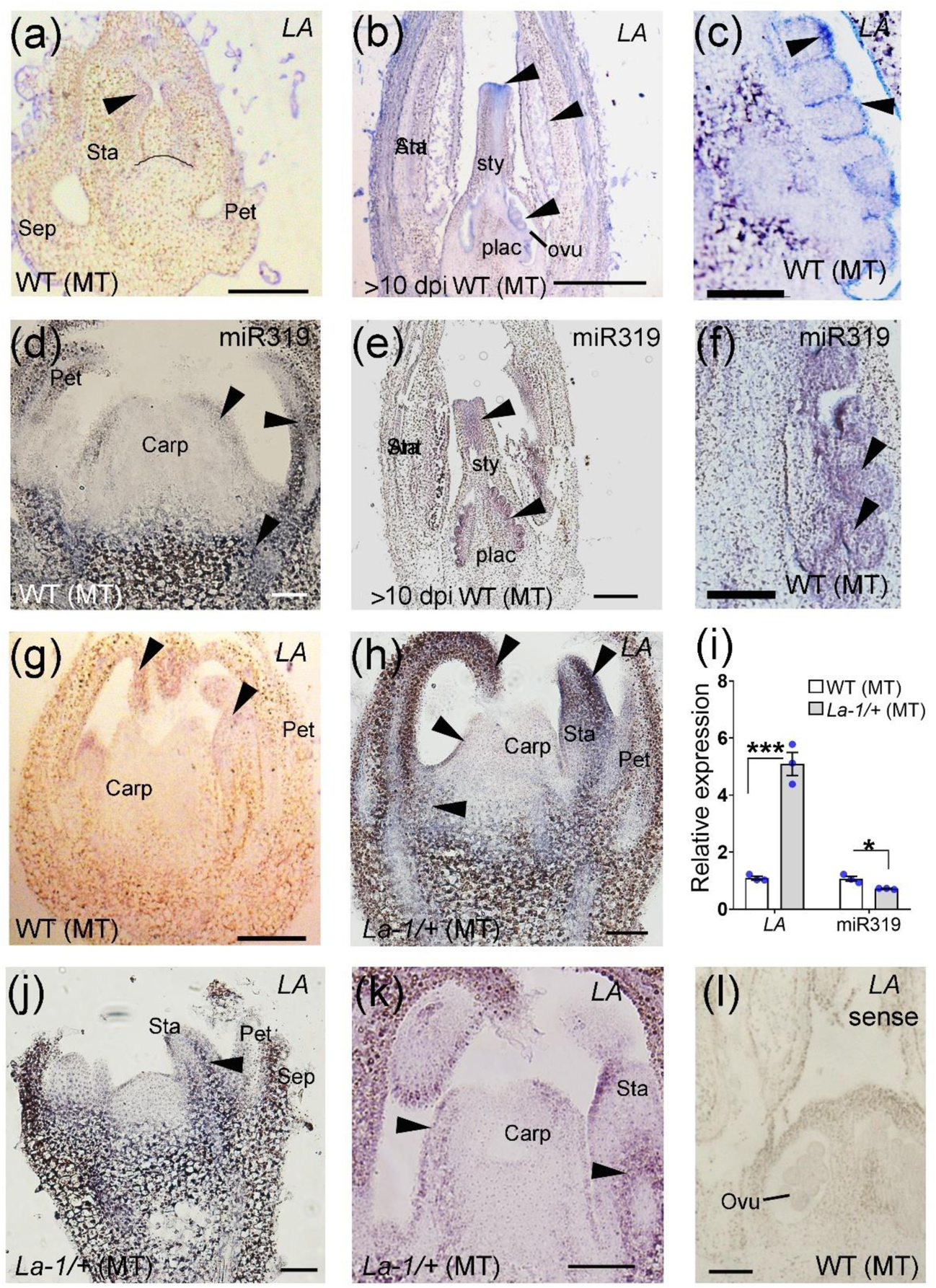
**Ectopic accumulation of *TCP4*/*LANCEOLATE* (*LA*) transcripts in *La-1/+* (MT) flowers.** (a-c) *TCP4/LA* transcripts detected by a digoxigenin labelled-*LA* antisense probe in WT (MT) 5-(days post-inflorescence) dpi flower bud (a), mature (>10 dpi) flower (b), and developing ovules of mature flowers (c). Line in WT (MT) flower bud (a) indicates the position from where the carpel will arise. Arrowheads indicate *LA* expression pattern. (d-f) miR319 transcripts detected by a digoxigenin labelled-*miR319* antisense probe in WT (MT) flower bud (d), mature flower (e), and developing ovules of mature flowers (f). Arrowheads indicate miR319 expression pattern. While *TCP4/LA* transcripts were detected only in petals and anthers of WT (MT) developing flower buds (g), they accumulate broadly in developing flower buds of *La-1/+* (MT) plants (h). (i) Relative expression levels of *TCP4*/*LA* and miR319 in 5-8 dpi floral buds from WT (MT) and *La-1/+* (MT) (n =3). \**P*<0.05 according to Student’s t-test (two-tailed). Values are means ± s.e. (j, k) Ectopic *TCP4/LA* transcript accumulation in very early stages of gynoecium development (j), mainly on the abaxial side of developing carpels (k). (l) No signal is detected by using a digoxigenin labelled-*LA* sense probe in WT (MT) flower bud. Sep: sepal; Pet: petal; Sta: Stamen; Sty: style; Carp: carpel; Plac: placenta; Ovu: ovule. Scale bars: 100 µm.

On the other hand, miR319 transcripts accumulated in most tissues of the developing ovules (Figure 3f, arrowheads), and are also localized in the anthers, stigma, and style at later stages (P2) of gynoecium development (Figure 3e). At P1, miR319 was expressed in petals, and in the margins and base of the developing carpel (Figure 3d). Given that *LA* is lowly expressed in early stages of carpel development (Figure 3a, g), we predicted that the precocious differentiation of the *La-1/+* carpel is a result of early *LANCEOLATE* de-repression. Indeed, *La-1/+* (MT) carpels at P1 already showed a strong *LA* expression in petal/sepal primordia, as well as in the innermost organs as anther and carpel primordia (Figure 3h), which partially overlaps with miR319 expression (Figure 3d). Accordingly, *LA* transcripts accumulated at higher levels in *La-1/+* (MT) reproductive meristem (Silva *et al*., 2019) and 5-8 dpi young flower buds (Figure 3i). miR319 transcripts, on the other hand, are reduced in 5-8 dpi *La-1/+* buds, suggesting a negative feedback regulation of the miR319/*LA* module. *LA* was ectopically expressed at very early stages of bud development in *La-1/+* (MT) plants (Figure 3j), when comparing with similar stages of WT (MT) plants (Figure 3a). Importantly, *LA* transcripts became localized in the abaxial margins and vasculature of the developing carpel primordia, as well as strongly expressed in the anther primordium of *La-1/+* (MT) young flower buds (Figure 3k). No signals were seen either with a control *LA* sense probe (Figure 3l) or a scrambled miRNA probe (not shown). Altogether, these observations suggest a tuning model for the interplay between *LA* and miR319, in which miR319 temporally down-regulates *LA* activity during early gynoecium patterning.

In WT (MT) gynoecia, *TCP4*/*LANCEOLATE* transcripts were lowly or even not detected in the carpel primordia tissues (Figure 3). Therefore, we predicted that reduced *LA* expression in tomato young flower buds is a prerequisite for the maintenance of carpel morphogenetic competence. We then evaluated flower and carpel phenotypes in *la-6* (M82) mutant plants, which contain a nonsense mutation that suppresses the *La/+* gain-of-function phenotype in a dominant manner (Ori *et al*., 2007). Accordingly, the *la-6* (M82) mutant showed nearly normal carpel and fruit development (Figure S4). To check the possibility that other miR319-regulated *CIN-TCP* family members act redundantly with *LA*, we inspected flower, carpel, and fruit phenotypes of miR319-overexpressing M82 plants (Ori *et al*., 2007). Like *la-6* (M82), miR319 overexpressors did not show any obvious defects in carpel or fruit development (Figure S4). On the other hand, *la-6* (M82) mutant and miR319 overexpressor plants did show larger sepals, consistent with the *TCP4*/*LANCEOLATE* expression in this floral organ (Figure 3). Thus, the reason *la-6* (M82) mutant and the miR319 overexpressors failed to show strong carpel and fruit defects is because *TCP4*/*LANCEOLATE* (and likely other *CIN-TCPs*) is post-transcriptionally regulated by miR319 in WT (Ori *et al*., 2007), thus preventing the activity of *CIN-TCPs* in early development of the carpel primordium.

*LA* transcripts were ready detected on the abaxial (outside) region of *La-1/+* (MT) developing carpels (Figure 3). To further investigate the impact of ectopic *LA* activity on carpel development, expression of a miR319-resistant allele of *LA* (*La-2*; Ori et al., 2007) was transactivated by the *FILAMENTOUS FLOWER* (*FIL*) promoter, which is expressed early in cell layers on the abaxial side of floral organs (Sawa et al., 1999; Figure S5a). *FILpro>>La-2* (M82) plants displayed elongated ovaries and fruits, with ovary and fruit shape indexes > 1.0 (Figure 4a-c), resembling *La-1/+* (MT) (Figure 1, S1). Ectopic *LA* expression may lead to the repression of particular targets, blocking cell division and promoting ovary and fruit elongation. To distinguish between the functions of LANCEOLATE as an activator or repressor in ovary and fruit development, we fused *La-2* ORF in-frame to the sequence encoding the C-terminal activation domain of VP16 (Sadowski *et al*., 1988), driven by the *FIL* promoter (*FILpro>>La-2-VP16* (M82) plants). We reasoned that in developmental contexts in which LA acts primarily as a repressor, this fusion would lead to the activation of downstream events (Sarvepalli and Nath, 2011). Expression of *FILpro>>La-2-VP16* partially reverted WT leaf phenotype comparing with *FILpro>>La-*2 (Figure S5b; Ori et al., 2007). Importantly, *FILpro>>La-2-VP16* (M82) plants showed partially rescued of WT (M82) ovary and fruit phenotypes (Figure 4a-c), suggesting that, similar to leaves, *LA* acts mostly as a repressor in carpel and fruit developmental contexts. Thus, ectopically expressing a miR319-resistant version of *LA* in early flower development is sufficient to modulate tomato fruit morphology.

**Figure 4.**
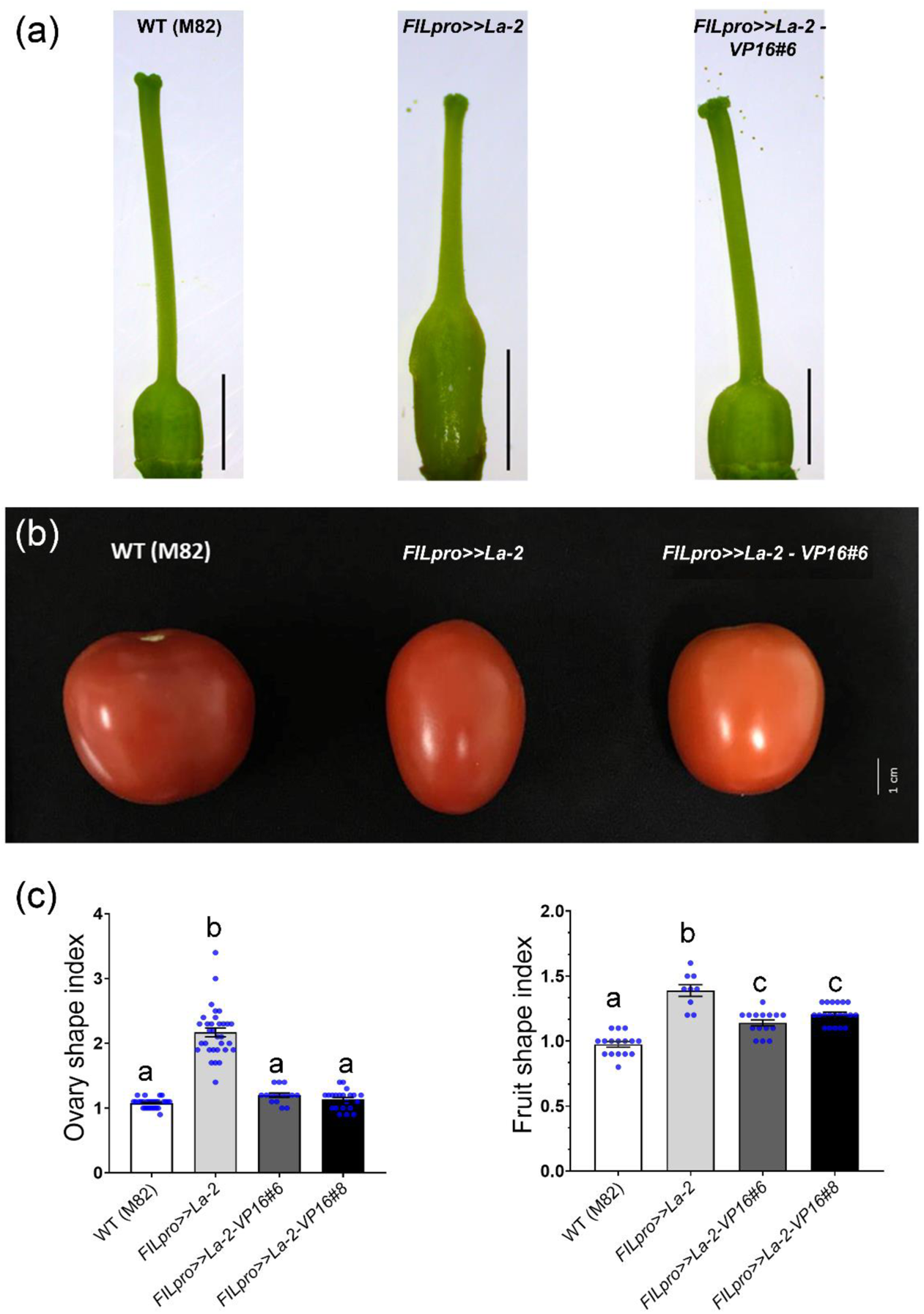
**miR319-targeted *TCP4/LA* acts mostly as a transcriptional repressor in carpel and fruit developmental contexts.** (a) Representative pre-anthesis carpels of WT (M82), *FILAMENTOUS FLOWER* promoter-driven *La-2* allele (*FILpro>>La-2*; Ori et al., 2007), and *FILpro>>La-2-VP16#6* line. Scale bars: 2 mm. (b) Representative fruits of WT (M82), *FILpro>>La-2*, and *FILpro>>La-2-VP16#6* line. Scale bar: 1 cm. (c) Ovary and fruit shape indexes of WT (M82), *FILpro>>La-2*, *FILpro>>La-2-VP16#6* and *FILpro>>La-2-VP16#8* lines. Values are mean ± s.e. Letters indicate the significant differences among different genotypes evaluated by Tukey’s HSD test (*P*< 0.05).

### *OVATE*, a major fruit shape gene associated with tomato domestication, is a direct target of LANCEOLATE

Fruit development in tomato and pepper is controlled by many Quantitative Trait Loci (QTL) but only a few have major effects on fruit shape variance, including *SUN* and *OVATE* (Gonzalo & Van Der Knaap, 2008; Xiao *et al*., 2009; Tsaballa *et al*., 2011). *OVATE* is highly expressed in young flowers buds (which include both P1 and P2 developmental stages; Huang et al., 2013), and it showed a opposite expression profile when comparing with *LA* during early flower development (Figure S3). Accordingly, *OVATE* was strongly down-regulated in 5-8 dpi *La-1/+* (MT) carpels (Figure 5a). Similarly, *ovate* null allele (Liu et al., 2002) introgressed into MT cultivar leads to an elongated fruit phenotype (Figure S6a). Such fruit phenotype is due to modifications in early ovary development in *ovate* (MT) flowers and reduced number of locules in the fruits (Figure S6b, c), resembling phenotypes observed in *La-1/+* (MT) (Figure 1). Recently, a modifier locus of *OVATE* was identified as another member of the OFP class, *SlOFP20* (Wu *et al*., 2018). In contrast to *OVATE*, the *SlOFP20* expression was not significantly altered in *La-1/+* (MT) floral buds (Figure 5a). This observation suggests that LANCEOLATE may affect gynoecium development partially by directly regulating *OVATE* expression. In addition, neither *LANCEOLATE* nor miR319 expression were significantly altered in developing gynoecia of *ovate* (MT) flowers (Figure S6d).

**Figure 5.**
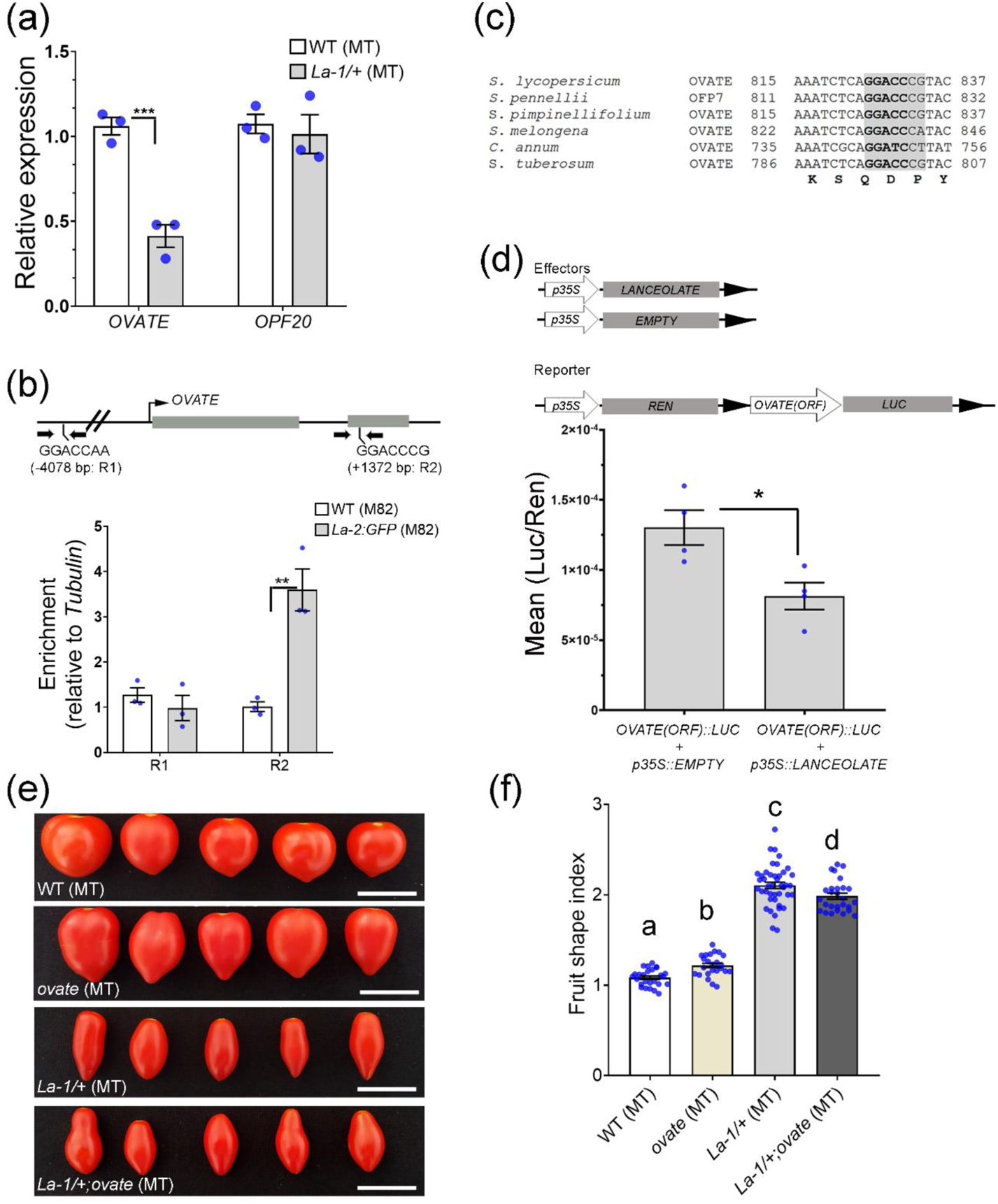
**The tomato domestication fruit shape *OVATE* is directly regulated by *TCP4/LANCEOLATE*.** (a) Relative expression levels of *OVATE* and *OPF20* in 5-8 days-post inflorescence (dpi) buds of WT (MT) and *La-1/+* (MT) (n =3). \*\*\**P*<0.001 according to Student’s t-test (two-tailed). Values are means ± s.e. (b) Upper panel, schematics of *OVATE* locus showing the genomic regions tested by Chromatin Immunoprecipitation (ChIP)-qPCR. Arrows indicate location of the specific oligos. Gray boxes, exons. Lower panel, TCP4/LA binding in the *OVATE* locus confirmed by ChIP-qPCR at the second exon (+1372 bp). \*\**P*<0.01, according to Student’s t-test (two-tailed). Values are mean ± s.e (n=3). R1 (putative TCP4/LA binding motif at −4078 bp), R2 (putative TCP4/LA binding motif at +1372 bp). *La-2:GFP*, *LApro>>La-2:GFP* (M82) plants. (c) Multiple sequence alignment of *OVATE-like* nucleotide sequences. *S. lycopersicum*, Solyc02g08550; *S. pennellii*, Sopen02g030180; *S. pimpinellifolium*, Sopim02g085500; *S. melongena*, SMEL_000g030240; *C. annum*, CA02g22830; *S. tuberosum*, PGSC0003DMP400022490. (d) Dual-luciferase transient expression assay (n=4) (\**P*<0.05 according to Student’s t-test two-tailed. Values are mean ± s.e). (e) Representative fruits of WT (MT), *ovate* (MT), *La-1/+* (MT), and double mutant *La-1/+;ovate* (MT). Scale bars = 2 cm. (f) Fruit shape index is the ratio of length to width. Values are mean ± s.e. Letters indicate the significant differences among different genotypes evaluated by Tukey’s HSD test (*P*< 0.05).

Screening for conserved *TCP4-like* consensus binding sites (Kosugi & Ohashi, 2002) revealed two partially conserved core motifs in the *OVATE* locus (Supplementary Table S1): one around 4000 bp upstream the start codon, and another in the second exon (Figure 5b). To validate the binding of the LANCEOLATE protein to the *OVATE* locus, we performed Chromatin Immunoprecipitation (ChIP)–qPCR experiments of plants carrying a *LApro>>La-2:GFP* (green fluorescent protein) transgene using young flower buds as a sample. *LApro>>La-2:GFP* (M82) plants express a miR319-resistant *LA* allele (Burko *et al*., 2013) and display modified pre-anthesis ovaries as well as flowers with shorter sepals (Figure S2e). We found a significant enrichment of La-2–GFP protein bound to the GGACCCG motif in the second exon (Figure 5b), consistent with the direct regulation of *OVATE* by LANCEOLATE.

TF binding sites located in exons are termed “duons” because they could influence the regulation of both transcription and amino acid sequence. Therefore, general synonymous nucleotide mutations should be under relaxed purifying selection in duons, because of the transcription factor recognition (Burgess *et al*., 2019). Accordingly, the LANCEOLATE binding site and adjacent regions are highly conserved among Solanaceae *OVATE-like* genes (Figure 5c). To provide additional evidences that LA can directly repress *OVATE* expression, we performed luciferase transient expression assays in *Nicotiana benthamiana* leaves. Our data showed that LA significantly repressed the *OVATE* exon sequence-driven luciferase activity (Figure 5d), reinforcing the direct regulation of *OVATE* by LANCEOLATE.

To further elucidate the interplay between *LA* and *ovate*, we introduced the *ovate* null allele (Gonzalo & Van Der Knaap, 2008) into *La-1/+* (MT) plants to generate *La-1/+*;*ovate* (MT) plants. *La-1/+*;*ovate* (MT) plants displayed simpler leaves and vegetative architecture similarly to *La-1/+* (MT) plants (Figure S6e). Most importantly, *La-1/+*;*ovate* (MT) ovaries at anthesis and fruits resemble those of *La-1/+* (MT) plants, but with slightly reduced fruit shape index (Figure S6f, g; Figure 5e and 5f). Together, our findings indicated that *LANCEOLATE* is epistatic and directly regulates a major fruit shape gene associated with tomato domestication, but *LANCEOLATE* and *OVATE* can also act in parallel pathways during fruit development.

### *LANCEOLATE* modulates auxin homeostasis in gynoecium development by directly regulating *SlYUCCA4*

Variation in tomato fruit morphology results from the action of fruit shape-associated genes in coordination with phytohormones, mainly auxin, at early stages of flower development (Seymour *et al*., 2013; Wang *et al*., 2019). TCP-mediated cell differentiation and organ maturation has been linked to the biosynthesis and response to auxin (Lucero *et al*., 2015; Koyama *et al*., 2010). Therefore, in addition to the *OVATE* direct repression by LANCEOLATE (Figure 5), the strong medial constriction in the *La-1/+* (MT) ovaries and fruits (Figure 1) may be due to modified responses to auxin. To investigate the possible link between auxin and *La-1/+* (MT) fruit development, we crossed transgenic MT plants carrying the auxin marker *pDR5::GUS* with *La-1/+* (MT) plants, generating *La-1/+* (MT); *pDR5::GUS* plants. At 4 dpi (Xiao *et al*., 2009), GUS staining was observed in petal and stamen primordia in MT; *pDR5::GUS* flower buds, but not in *La-1/+* (MT); *pDR5::GUS* flower buds (Figure 6a, f). During the onset of carpel primordia (5-6 dpi; Xiao *et al*., 2009), GUS staining was detected in the tip of carpel and stamen primordia in MT; *pDR5::GUS*, but only in stamen primordia of *La-1/+* (MT); *pDR5::GUS* (Figure 6b, g). At 7-8 dpi, *GUS* expression marks the region of further placental development in MT; *pDR5::GUS*, while GUS staining is more prominent in the tip of unfused and fused carpel primordia of *La-1/+* (MT); *pDR5::GUS* buds (Figure 6c, d, h, i). These results suggested that auxin response is lower at very early stages of *La-1/+* (MT) gynoecium patterning. After 10 dpi, auxin response becomes localized in the ovules of MT; *pDR5::GUS* flowers, but rather localized at the base of *La-1/+* (MT); *pDR5::GUS* ovaries (Figure 6e, j).

**Figure 6.**
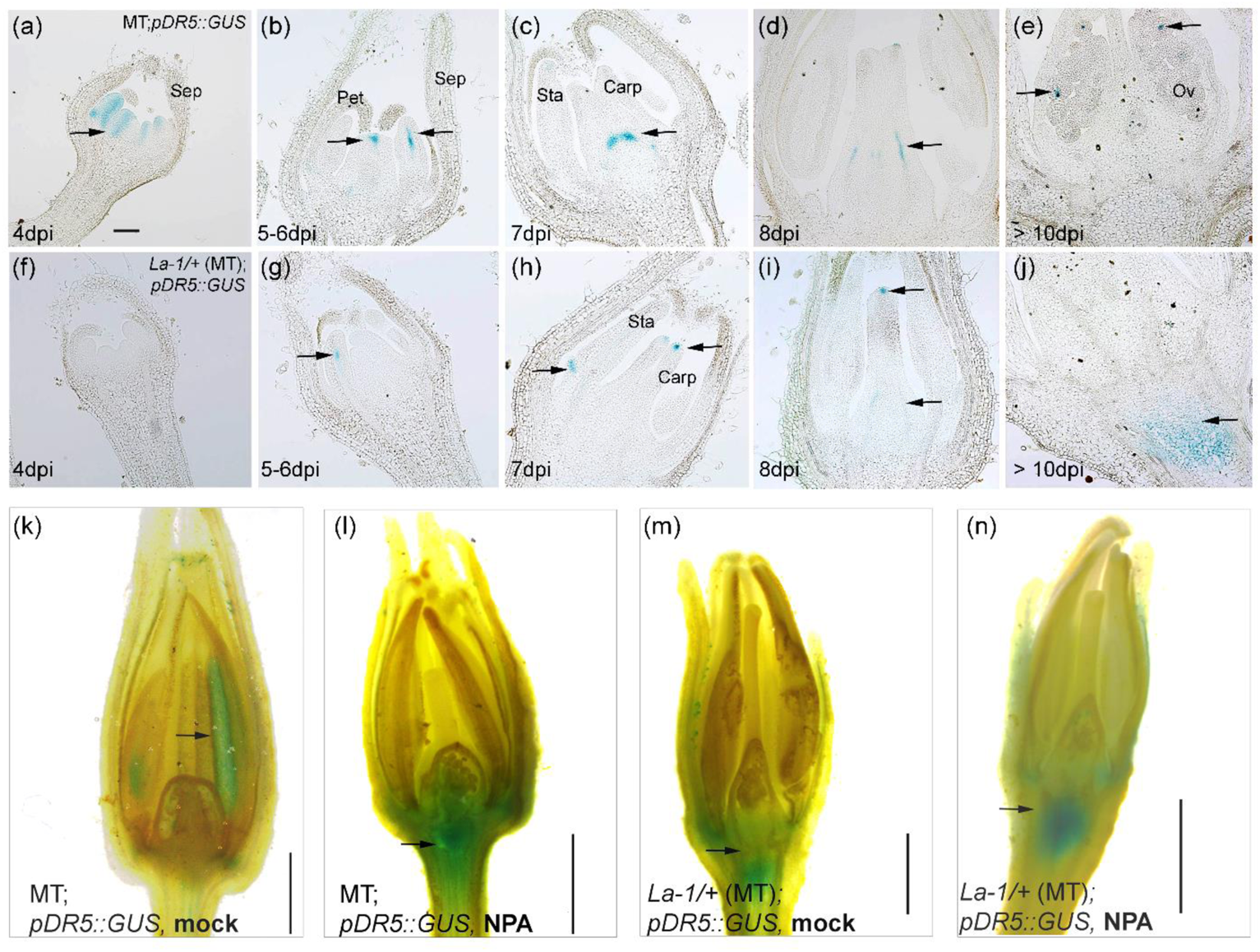
**MiR319-targeted *TCP4*/*LANCEOLATE* de-repression leads to alteration is auxin responses during early gynoecium patterning.** (a-j) Auxin reporter gene assay by GUS staining (arrows) of WT (MT) (a-e) and *La-1/+* (WT) floral buds (f-j) at different days post-inflorescence (dpi). Bar = 100 µm. (k-n) Auxin transport inhibitor *N*-1-naphthylphthalamic acid (NPA) treatment of 12-dpi floral buds of MT; *pDR5::GUS* and *La-1/+*(MT); *pDR5::GUS* plants. Arrows indicate GUS staining after mock or NPA treatments. Scale bars = 1 mm (k, m), 2 mm (l, n).

Our data suggest that a fine-tunning auxin response at early developmental stages of the gynoecium is crucial for proper fruit development. Interestingly, tomato fruits treated with the auxin transport/response inhibitor *N*-1-naphthylphthalamic acid (NPA) show larger pedicels and an increase in *pDR5::GUS* expression in the base of the ovary, placenta, and columella (Pattison & Catalá, 2012a). *La-1/+* (MT) fruit pedicels showed a distal/proximal ratio significantly (*P* < 0.001) greater than WT (MT) pedicels (Figure S7). This indicated that pedicel growth was stimulated in *La-1/+* (MT) plants, similarly to NPA-treated tomato and apple flowers (Drazěta *et al*.; Pattison & Catalá, 2012b). Given that *LANCEOLATE* de-repression modifies auxin response (Figure 6a-j), we treated 12-dpi flowers from MT; *pDR5::GUS* and *La-1/+* (MT); *pDR5::GUS* with NPA for 2 days. As expected, NPA treatment caused a substantial increase in *pDR5::GUS* expression in MT flowers, mostly in the placenta and the base of the ovary (Figure 6l), when comparing with mock-treatment in which GUS staining is observed only in the stamens (Figure 6k). On the other hand, *pDR5::GUS* expression did not substantially changed in NPA-treated *La-1/+* (MT); *pDR5::GUS* flowers when comparing with mock-treated 12-dpi flowers (Figure 6m, n). In addition, *GUS* expression in *La-1/+* (MT); *pDR5::GUS* flowers is detected in the base of ovary, similarly to MT; *pDR5::GUS* flowers treated with NPA (Figure 6l-n). These findings reinforce the idea that auxin response is modified at early stages of in *La-1/+* (MT); *pDR5::GUS* flower development. Thus, we concluded that *LANCEOLATE* de-repression in reproductive apices led to altered auxin response in gynoecium.

We then hypothesized that *LANCEOLATE* might regulate auxin responses via modulating auxin transport in the carpels. PIN-FORMED (PIN) proteins are auxin efflux facilitators (Friml *et al*., 2003). *AtPIN1* is the most important member of the *Arabidopsis* PIN family, and the single *pin1* mutant exhibits severe flower developmental defects (Okada *et al*., 1991). Given that *AtPIN1* is the orthologue of tomato *PIN1* (*SlPIN1*) and that GFP localization in tomato plants harbouring *pAtPIN1::PIN1-GFP* is similar at vegetative and reproductive organs when comparing with endogenous tomato PIN1 localization (Bayer *et al*., 2009; Goldental-Cohen *et al*., 2017), we crossed transgenic MT plants carrying *pAtPIN1::PIN1-GFP* with *La-1/+* (MT) to generate *La-1/+* (MT); *pAtPIN1::PIN1-GFP* plants. However, we did not see any obvious modifications in PIN1-GFP protein accumulation or localization in developing carpels of *La-1/+* (MT); *pAtPIN1::PIN1-GFP* when comparing with MT; *pAtPIN1::PIN1-GFP* carpels at similar developmental stages (Figure S8). As previously reported, PIN1-GFP proteins were localized toward the tip of the gynoecium, but they were also present in brefeldin A (BFA) body-like (Goldental-Cohen *et al*., 2017). These observations indicate that the modified auxin response in *La-1/+* ovaries (Figure 6) is not a result of modifications in the PIN1-dependent auxin transport.

To further investigate the possible molecular mechanisms underlying auxin responses in *La-1/+* (MT) flowers, we examined several genes associated with auxin metabolism, signal transduction and homeostasis, which have been reported to be expressed in tomato early flower development (Expósito-Rodríguez *et al*., 2011; Wang *et al*., 2019; Wu *et al*., 2011; Wu *et al*., 2012; Zouine *et al*., 2014). While *SlYUCCA1* and *-6* expression did not change, *SlYUCCA3* and *SlYUCCA4* were up- and down-regulated, respectively, in 5-8 dpi *La-1/+* (MT) buds (Figure 7a). In terms of signal transduction, *SlARF10B* was down-regulated in the *La-1/+* (MT) young buds, while transcript levels of *SlIAA14*, *SlARF3*, *SlARF10A*, *SlARF16A*, and *SlARF17* were unaltered (Figure 7b, d). The *Gretchen Hagen 3* (*GH3*) genes encode acyl acid amido synthetases, many of which have been shown to modulate the amount of active auxin, including *SlGH3.4* and *SlGH3.9* (Wang et al., 2019). The down-regulation of *SlGH3.9* in *La-1/+* (MT) buds (Figure 7c) may lead to altered auxin homeostasis early in flower development.

**Figure 7.**
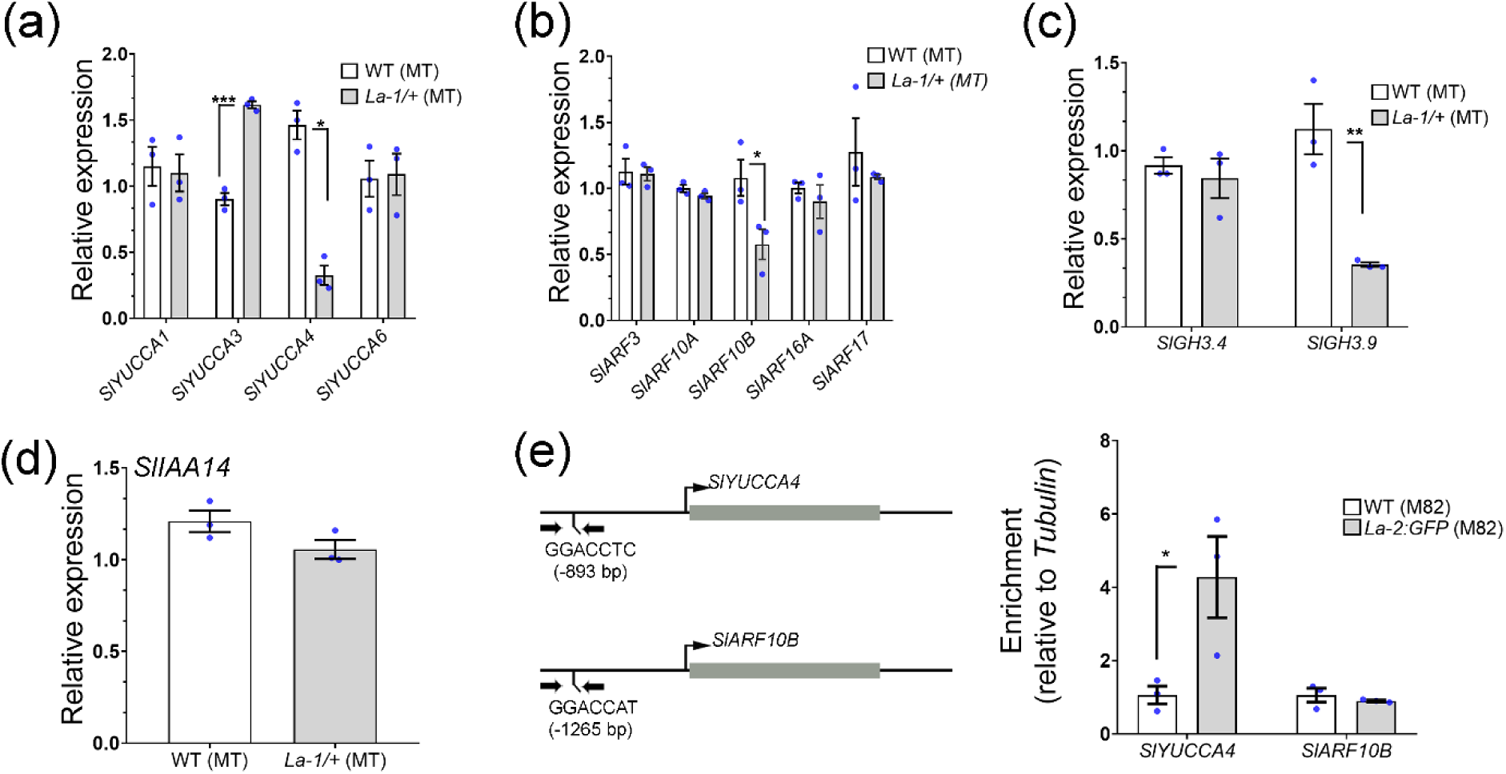
**TCP4/LA regulates auxin responses in floral buds by directly binding to *SlYUCCA4* promoter.** (a) Relative expression levels of *SlYUCCA1, -3, -4,* and *-6* in 5-8 days-post inflorescence (dpi) buds of WT (MT) and *La-1/+* (MT) (n =3). (b) Relative expression levels of *SlARF3, SlARF10A, SlARF10B, SlARF16A,* and *SlARF17* in 5-8 days-post inflorescence (dpi) buds of WT (MT) and *La-1/+* (MT) (n =3). (c) Relative expression levels of *SlGH3.4* and *SlGH3.9* in 5-8 days-post inflorescence (dpi) buds of WT (MT) and *La-1/+* (MT) (n =3). Relative expression level of *SlIAA14* in 5-8 days-post inflorescence (dpi) buds of WT (MT) and *La-1/+* (MT) (n =3). \**P*<0.05, \*\**P*<0.01, \*\*\**P*<0.001 according to Student’s t-test (two-tailed). Values are means ± s.e. (e) Left panel, schematics of *SlYUCCA4* and *SlARF10B* loci showing the genomic regions tested by Chromatin Immunoprecipitation (ChIP)-qPCR. Arrows indicate location of the specific oligos. Gray boxes, exons. Right panel, TCP4/LA binding in the *SlYUCCA4* and *SlARF10B* loci by ChIP-qPCR. \**P*<0.05, according to Student’s t-test (two-tailed). Values are mean ± s.e (n=3). *La-2:GFP*, *LApro>>La-2:GFP* (M82) plants.

In *Arabidopsis*, the LANCEOLATE homologue TCP4, controls auxin biosynthesis by directly regulating *YUCCA5* (Challa *et al*., 2016). We then screened for putative conserved *TCP4-like* binding sites in the misexpressed genes *SlYUCCA3, SlYUCCA4*, *SlARF10B*, and *SlGH3.9* (Figure 7a-d). We found one site (GGACCTC) in the *SlYUCCA4* promoter (localized at −893 bp from the start codon), and one site (GGACCAT) in the *SlARF10B* promoter (localized at −1265 bp from the start codon; Supplementary Table S1). Only *SlYUCCA4* promoter showed enrichment for the binding of the La-2–GFP protein (Figure 7e). Taken together, our data indicated that *LANCEOLATE* controls auxin biosynthesis during tomato flower development by directly repressing *SlYUCCA4*.

Although it is not clear how *OVATE* affects fruit development via auxin response, some genes associated with auxin biosynthesis, signal transduction and homeostasis were reported to be misexpressed in gynoecium of *S. pimpinellifolium* carrying an *ovate* mutation (Wang *et al*., 2019). We then tested whether auxin-associated genes misregulated in *La-1/+* (MT) (Figure 7) were also misexpressed in *ovate* (MT) buds. *SlYUCCA3* and *SlYUCCA4* were up- and down-regulated, respectively, in *ovate* (MT) (Figure S9a), similarly to *La-1/+* (MT) buds (Figure 7). On the other hand, *SlARF10B* and *SlARF16A* transcript levels slightly increased in 5-8 dpi *ovate* (MT) flowers, when comparing with WT (MT) (Figure S9b). Likewise, *SlIAA14* and *SlGH3.9* were up-regulated in *ovate* (MT) buds (Figure S9c, d). Our observations suggest that *LANCEOLATE* and *OVATE* similarly modulate auxin biosynthesis in flower buds, but auxin signal transduction and homeostasis are distinctly affected by these two frui morphology-associated factors. This may explain the more constrict and elongated ovaries and fruits observed in *La-1/+* (MT) plants when comparing with *ovate* (MT) (Figure 5).

### *Arabidopsis* miR319-targeted *CIN-TCPs* also modulate auxin homeostasis in developing gynoecia

The *Arabidopsis tcp4-soj8* mutant contains a point mutation in the miRNA target site of the *LANCEOLATE* homologue, *TCP4*, that lessens the interaction with miR319, thus de-repressing *TCP4* (Palatnik *et al*., 2007). To investigate the possible roles of *TCP4* and other *CIN-TCPs* in *Arabidopsis* fruit development, we compared fruit phenotype of Columbia (Col-0) wild-type and *tcp4-soj8*, *Jaw-D*, and *tcp2-3-4-10* mutants (Bresso *et al*., 2018). While *tcp4-soj8* siliques were smaller than Col-0, *Jaw-D* and *tcp2-3-4-10* mutants (Nag *et al*., 2009; Bresso *et al*., 2018) showed siliques with crinkled surface (Figure S10a). Similar to tomato (Figure 1), the miR319-dependent *CIN-TCP* regulation is critical for proper *Arabidopsis* gynoecium development. Similar to crinkled leaves, the crinkled fruit phenotype might be a result of altered auxin responses (Lucero *et al*., 2015). Given that auxin homeostasis is altered in developing gynoecium of *La-1/+* (MT) flowers (Figure 5), we crossed Col-0; *pAtPIN1::PIN1-GFP* with *tcp4-soj8* and *Jaw-D* mutants. PIN1-GFP accumulated similarly in the developing gynoecia of all genotypes, indicating that *PIN1* expression was not perturbed by *Arabidopsis CIN-TCP* mis-regulation in inflorescences at developmental stages 1 to 7 (Figure S10b). We then measured the expression of the auxin biosynthesis genes *YUCCA1* and *-4* in inflorescences (developmental stages 1 to 7) of Col-0, *tcp4-soj8*, *tcp2-4, tcp2-3-4-10,* and *Jaw-D*. Although we did not observed a significantly alteration in *tcp2*-*4* double mutant, *YUCCA1* and *-4* were significantly up-regulated in the quadruple mutant *tcp2-3-4-10* and *Jaw-D* developing gynoecia (Figure S10c). Together, our data suggest that auxin biosynthesis is redundantly regulated by miR319-targeted *CIN-TCPs* during *Arabidopsis* gynoecia development, similarly to what was described for the class I *TCP15* (Lucero *et al*., 2015). Importantly, the negative regulation of auxin biosynthesis at early stages of gynoecium development is conserved between tomato *LANCEOLATE* and *Arabidopsis CIN-TCPs*.

*OVATE* is a direct target of *LANCEOLATE* in tomato (Figure 5) and the dominant *Atofp1-1D* mutant displays reduced silique length caused by reduced cell elongation (Wang *et al*., 2007). However, *OFP1* locus does not contain conserved *CIN-TCP* binding sites (Kosugi & Ohashi, 2002) (not shown), and it was similarly expressed in Col-0, *tcp4-soj8*, and *Jaw-D* developing gynoecia (Figure S10d). Thus, the function of *Arabidopsis CIN-TCPs* in gynoecium patterning and fruit development seems to be independent of *OFP1* regulation.

## DISCUSSION

Despite many studies describing the roles of *CIN-TCPs* on vegetative development, surprisingly little is known about the molecular mechanisms by which *CIN-TCPs* regulate reproductive development, mainly carpel and fruit development (Crawford *et al*., 2004; Nag *et al*., 2009; Lucero *et al*., 2015; Silva *et al*., 2019). Indeed, only *Arabidopsis TCP14* and *-15* were known to regulate gynoecium patterning (Lucero *et al*., 2015). Here, we show that the tomato TCP4 homologue, LANCEOLATE, plays an important role in the final fruit morphology and development (Figure 1). We found the miR319 regulation at early stages of tomato flower development limits *TCP4*/*LANCEOLATE* (and perhaps other *CIN-TCPs*) function during morphogenesis and maturation of flower whorls. We highlighted that miR319 regulation is crucial for carpel primordia growth until mature carpel, especially in the outer layer of floral meristem and later in the stigma and outermost layer of the ovules (Figure 3). The *La-1/+* (MT) mutant flower buds present an ectopic *TCP4*/*LANCEOLATE* expression in the inner layers of floral meristem and carpel primordia (Figure 3h, j-k), where it represses tissue growth likely by negatively modulating cell division. Similarly, *TCP4*/*LANCEOLATE* spatiotemporal expression is required at the marginal blastozone of developing leaf for maintenance of leaf morphogenetic window and proper compound leaf formation (Ori et al., 2007; Shleizer-Burko et al., 2011). Hence, the ectopic expression of *TCP4*/*LANCEOLATE* in the inner tissue of developing *La-1/+* (MT) carpel primordia is likely associated with the absence of placenta and reduced ovule formation due to premature maturation of these tissues.

Indeed, the limiting spatiotemporal expression of *CIN-TCPs* during flower development seems to be a prerequisite for proper reproductive development in different species. *Arabidopsis* plants with higher and ectopic CIN-TCP activity in flowers exhibits absence of petals and stamens, and modified carpels (Nag *et al*., 2009). *CIN-TCP* mis-expression in *Antirrhinum* flowers negatively affects petal growth (Crawford *et al*., 2004). Our data indicate that, in both tomato and *Arabidopsis*, the miR319-*CIN-TCP* module plays a role in leaf and carpel development (Figure 1, S10). This dual role in leaf and carpel development is not restricted to the miR319-*CIN-TCP* module. In tomato, for instance, the loss of miR156-targeted *SPL/SBP* function leads to smaller and simpler leaves, and alterations in the maintenance of the meristematic state of carpel tissues (Silva *et al*., 2014). These miRNA-associated regulatory modules were likely co-opted for the regulation of gynoecium development.

We propose a model in which the spatiotemporal fine-tuning of *TCP4*/*LANCEOLATE* expression by miR319 determines the cell growth patterns underlying tomato fruit morphology mostly via two points: (i) directly repressing *OVATE*, a negative regulator of fruit elongation; and (ii) directly repressing *SlYUCCA4*, thereby reducing auxin response (Figure 8). The discovery that *OVATE* is a direct target of TCP4/LANCEOLATE was surprising, considering that *OVATE* is a major QTL associated with domestication events that led to tomato fruit shape diversity (Liu *et al*., 2002; Rodriguez *et al*., 2011). Therefore, our work offers the first molecular link between a microRNA module and a domestication-associated gene during the cell proliferation and differentiation balance in tomato gynoecium patterning. Interestingly, this molecular link seems to be not conserved in a dry fruit-bearing species (Figure S10), which might be at least one of the reasons of why the modulation of *LANCEOLATE* and *TCP4* expression, in tomato and *Arabidopsis*, respectively, leads to distinct fruit phenotypes. The role of miR319-*CIN-TCP* module in gynoecium patterning illustrates that the influence of a particular miRNA module on fruit shape depends on downstream targets and fruit type.

**Figure 8.**
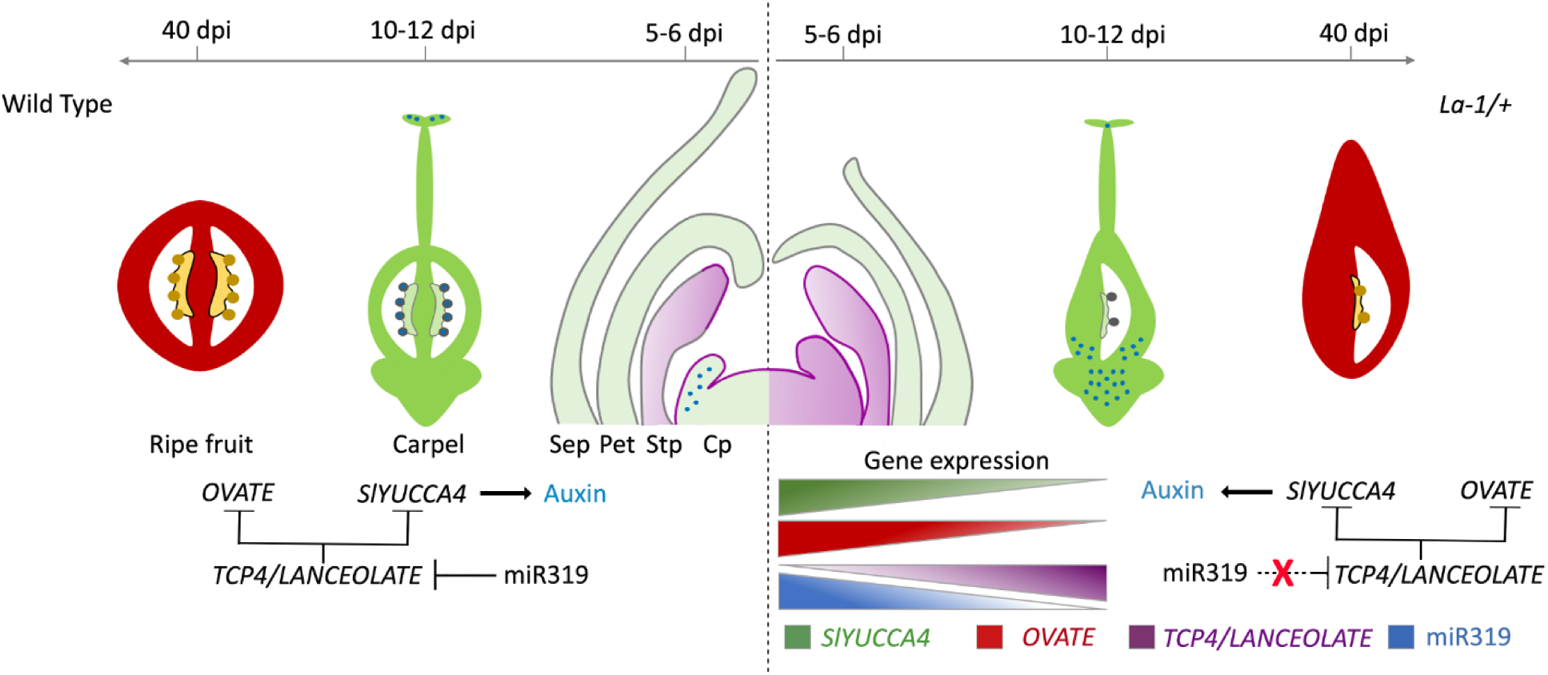
Proposed model for gynoecium patterning and fruit shape establishment orchestrated by the crosstalk between miR319-targeted *TCP4*/*LANCEOLATE (LA)*, the fruit shape *OVATE*, and auxin. The loss of miR319-dependent regulation of *LA* in the semi-dominant *Lanceolate* mutant (*La-1/+*) leads to ectopic *LA* expression in floral meristem and repression of *OVATE* expression in developing carpels. In WT (MT), auxin (light blue circles) is observed at early gynoecium patterning (5-12 days post-inflorescence or dpi) on the tip of the carpel, whereas *TCP4*/*LANCEOLATE* negatively modulates auxin metabolism by repressing *SlYUCCA4* in *La-1/+* (WT) gynoecia. Later in carpel development, auxin is localized in the base of *La-1/+* (WT) ovaries (light blue circles). By modulating auxin metabolism during flower maturation, the miR319/*LANCEOLATE* regulatory hub adjusts auxin responses to control cell proliferation and differentiation in tomato developing carpels. Black T-bar, direct repression effect; Purple, *LA* expression; Green, *SlYUCCA4* expression; Red, *OVATE* expression; Blue, miR319 expression. Sep: sepal; Pet: petal; Stp: stamen primordia; Cp: carpel primordia.

The strength of the fruit phenotypes due to *La-1* mutation varies among cultivars and accessions (Figure 1, S2), suggesting that interactions between *TCP4/LANCEOLATE* and uncharacterized genetic modifiers can also affect tomato final fruit shape. Similar phenotypic variation is observed for classical fruit shape-associated mutations. For instance, *OVATE* mutation leads to a range of fruit shapes from elongated and obovoid to round fruits, depending on the cultivars and accessions (Rodriguez *et al*., 2011). It will be interesting in near future to identify the possible modifier(s) that modulate *TCP4/LANCEOLATE* activity during gynoecium development and fruit set in different cultivars.

*TCP4/LANCEOLATE* and perhaps other miR319-targeted *CIN-TCPs* form a miRNA-dependent ‘regulatory hub’ that connects auxin to the fruit morphogenetic program in the control of cell growth and differentiation, which may explain the more extreme fruit shape alterations mediated by *TCP4/LA.* Local auxin production and signalling, along with the transport of auxin within the gynoecium is crucial for generation of the auxin gradient required for the apical-basal axis specification of *Arabidopsis* gynoecium (Robert *et al*., 2015a). Auxin dynamics is modified during *La-1/+* (MT) gynoecium patterning, at least in part by the direct repression of *SlYUCCA4* expression by TCP4/LA. As a result, *La-1/+* (MT) floral buds show lower auxin responses in early flower developmental stages, and premature auxin responses in the base of the ovaries (Figure 6). Moreover, *La-1/+* (MT) fruits show rudimentary placenta, and hence reduced ovule formation, which might be a result of lower auxin responses during gynoecium patterning. In fact, the expression of *MONOPTEROS* (*MP*/*ARF5*) in early stages of placenta development is crucial for ovule primordia formation, and *mp* mutants do not develop neither placenta nor ovules (Galbiati *et al*., 2013). Proper auxin dynamics throughout ovule and embryo development (Robert *et al*., 2015) is crucial for induction of the gibberellin biosynthesis required for pericarp growth after anthesis (Azzi *et al*., 2015). The lower number of cell layers in *La-1/+* (MT) fruit pericarp at 16 DPA (Figure 2) might be a result of low GA levels in *La-1/+* (MT) fruits caused by reduction in auxin biosynthesis in the embryos.

How *OVATE* controls auxin responses during tomato ovary and fruit development is not well understood (Wang et al., 2009). Our data indicate that key components of the auxin pathway in tomato are misregulated in both *La-1/+* (MT) and *ovate* (MT) floral buds (Figure 7, S9). However, only *SlYUCCA3* and *SlYUCCA4* showed similar expression patterns in *La-1/+* (MT) and *ovate* (MT) floral buds, which indicates that auxin metabolism is similarly modified by either de-repression of *TCP4/LA* or loss of *OVATE* activity. On the other hand, the expression of genes associated with auxin signalling and homeostasis showed an opposite trend in both mutants, which may explain the differences observed in their final fruit morphology and seed number (Figure 1, S6h). Therefore, it is reasonable to assume that different factors in the auxin pathway have distinct roles in remodeling organ shapes, and that responses to auxin may depend on the genetic background. Given the striking variation in fruit morphology among members of the Solanaceae family, fine-tuning regulation of gene expression by miRNA regulatory hubs coupled with modulation of hormone dynamics may be a common driver in the evolution of fruit-shape diversity.

## MATERIAL AND METHODS

### Plant materials and growth conditions

Tomato (*Solanum lycopersicum*) plants were grown in the greenhouse under natural day length conditions. *La-1/+* mutants were previously described (Silva *et al*., 2019). Near-isogenic line of the *ovate* mutant in tomato cv. Micro-Tom (MT) was obtained as described (Lombardi-Crestana et al, 2012). MT*; pAtPIN1::PIN1-GFP* and MT; *pDR5::GUS* transgenic plants were obtained by introducing the *pAtPIN1::PIN1-GFP* and *pDR5::GUS* constructs, respectively, into tomato cv. MT (Lombardi-Crestana et al, 2012). At least five transgenic events were obtained with similar expression patterns for each genotype, and one event was used for further analyses and crossings. *FILpro>>La-2* (M82)*; LApro>>La-2:GFP* (M82), *LApro >>AtMIR319a* (M82), and *la-6* (M82) plants were described elsewhere (Ori *et al*., 2007; Burko *et al*., 2013). Tomato flower buds and gynoecia were collected as previously described (Xiao *et al*., 2009). For generating the *FILpro>>La-2-VP16* (M82) plants, the *La-2* ORF (a miR319-resistant allele of *TCP4/LA*; Ori et al; 2007) was fused to the VP16 herpes simplex virus-derived transactivation domain, and the *La-2-VP16* construct was transactivated by the *FILAMENTOUS FLOWER* (*FIL*) promoter as described (Ori et al, 2007). The transgenic plants were generated as described (Ori et al, 2007). At least five transgenic lines were obtained, and two were used for further analyses.

*Arabidopsis thaliana* genotypes were in Columbia (Col-0) background and they were grown under long-day conditions (16 h: 8 h, light: dark) at 22°C. Transgenic *AtPIN1::PIN1-GFP* plants and the mutants *tcp4-soj8, tcp2-4, tcp2-3-4-10,* and *Jaw-D* were described previously (Palatnik *et al*., 2003; 2007; Bresso *et al*., 2018; Goldental-Cohen et al., 2017).

### RNA isolation and transcriptional analyses

Total RNA was extracted using TRIzol reagent (Thermo Fisher Scientific) according to manufacturer’s instructions. Briefly, 2 ug of total RNA were treated with TURBO DNA-free Kit (Thermo Fisher Scientific) to remove any residual genomic DNA. The cDNA synthesis and qPCR conditions were previously described (Silva *et al*., 2019). The qPCR reactions were performed in the StepOnePlus Real-Time PCR System with three biological samples and technical triplicates. Expression levels were calculated relative to the housekeeping tomato *ACTIN* (Expósito-Rodríguez *et al*., 2008) and *Arabidopsis PP2A* using the 2^−ΔΔCT^ method (TD, 2001). Oligos are listed in the Table S2.

### Floral meristem image analysis

Dissection of reproductive apices and modified pseudo-Schiff propidium iodide (mPS-PI) staining were made as described (Bencivenga *et al*., 2016; Serrano-Mislata *et al*., 2017). Custom Python scripts and Fiji macros based on a previous set of scripts (Bencivenga *et al*., 2016) were used to segment confocal image stacks, measure shoot meristem areas, and define the position of cells within the shoot meristem as described (Serrano-Mislata *et al*., 2017).

### Histological analysis and Scanning Electron Microscopy (SEM)

Preanthesis flowers, flowers at anthesis and fruit pericarps at 16 days post-anthesis (DPA) from WT (MT) and *La-1/+* (MT) plants were collected and fixed in Karnovsky solution (Karnovsky, 1965), using vacuum-infiltrated for 15 min. Samples were dehydrated in an increasing ethanol series (10–100%), and infiltrated using a HistoResin embedding kit (Leica, www.leica-microsystems.com), according to the manufacturer’s instructions. Tissue sections were obtained using a rotary microtome (Leica), stained with toluidine blue 0.05% (Sakai, 1973) and photographed in a Leica DM-LB microscope (Heidelberg, Germany), coupled with a Leica DFC310 camera (Wetzlar, Germany). Histological analysis was carried out in the ovary and pericarp central region, where the area and cell layer number of cells were measured using ImageJ software (http://rsbweb.nih.gov/ij/). For SEM, samples were fixed, mounted, and analyzed as described in Bharathan et al. (2002).

### Pedicel diameter

Pedicel diameter of distal and proximal regions of MT e La-1/+ fruits was determined using a digital caliper (Western Ferramentas, São Paulo, SP, Brazil). The measurements were performed in pedicels of fruits at three different maturity stages (mature green, breaker or red ripening).

### GUS staining

Closed floral buds and gynoecia at 4-12 dpi were collected, cut longitudinally and placed into GUS staining buffer [80mM sodium phosphate buffer, pH 7.0; 8mM EDTA; 0.4 mM potassium ferrocyanide; 0.05% Triton X-100; 0.8 mg ml-1 5-bromo-4-chloro-3-indolyl-β-D-glucoronide (X-Gluc); 20% methanol]. Samples were vacuum-infiltrated for 15 min and incubated in the dark at 37°C overnight (16 h). Following GUS staining, the samples were washed several times with 70% ethanol to extract chlorophyll and photographed using a Leica S8AP0 microscope set to 80X magnification coupled to a Leica DFC295 camera.

### NPA treatments

NPA-containing and mock solutions were applied on 12-dpi flower buds of MT; *pDR5::GUS* and *La-1/+* (MT); *pDR5::GUS* plants. After two days, the samples were collected and placed into GUS staining buffer. Flower buds were photographed using a Leica stereomicroscope (Germain). NPA stock solution (100 mM) was prepared in dimethyl sulfoxide (DMSO). Plants were treated with 10µL of 50 µM NPA solution (Pattison & Catalá, 2012a). Mock solution was prepared with an equivalent concentration of DMSO (0.05% v/v).

### *In situ* hybridization

Developing flower buds (5-12 dpi) from WT (MT) and *La-1/+* (MT) plants were used for *in situ* hybridization. Locked nucleic acid (LNA) probes with sequence complementary to AtmiR319a (5’-TTGGACTGAAGGGAGCTCCCT) and negative control (scrambled miR, 5’-GTGTAACACGTCTATACGCCCA-3’) were synthesized by Exiqon (http://www.exiqon.com/), and digoxigenin-labeled using a DIG oligonucleotide 3’end labeling kit (Roche Applied Science). Ten picomoles of each probe were used for each slide. Probes sense and antisense for *TCP4/LANCEOLATE* (Solyc07g062680) were used as described by Javelle and Timmermans (2012) at 0.5 ng .µL^−1^.kb^−1^. Hybridization and washing steps were performed at 55°C as described by Javelle and Timmermans (2012).

### Crossings

A crossing between *La-1/+* (MT) and *ovate* (MT) mutants was performed. The presence of the *La-1* mutation in the F2 offspring was identified by the simpler leaf phenotype (Silva et al., 2019). For the *ovate* mutation, a derived cleaved amplified polymorphic sequence marker was developed (Table S2). After amplification, the PCR products were digested with *BsiWI* restriction enzyme and analysed in 1.5 % agarose gel. The tomato F1 hybrid offspring shown in the Fig. S2 was generated by crossing the semi dominant *La-1/+* mutant in LA0335 background with Micro-Tom (MT) and M82 cultivars.

### Chromatin immunoprecipitation (ChIP)

ChIP assay was performed on 5-10-days post floral initiation (dpi) flower buds from *LApro>>La-2:GFP* (M82) plants as described (Burko *et al*., 2013; Frank *et al*., 2018), with few modifications. Briefly, 1.5 g of flower tissues were cross-linked 15 min in vacuum infiltration with 1.0 % v/v formaldehyde solution. Subsequently, nuclei were isolated and chromatin was fragmented by sonication to obtain DNA fragments ranging between 0.3 – 1 Kb. Immunoprecipitation was performed using EpiQuik Plant ChIP kit (Epigentek Group, USA) following manufacturer’s instructions. DNA-protein complexes were immunoprecipitated using a monoclonal anti-GFP antibody (Roche). Analysis of enrichment of target genes was performed by qPCR using the oligos listed in the Table S2. Enrichment for TCP4/LANCEOLATE-bound sequences was assayed by quantitative PCR on the immunoprecipitated DNA. Quantitative PCR enrichment was calculated by normalizing to *TUBULIN* and to the total input of each sample. Two technical repeats and three biological replicates per genotype were analysed for the enrichment of each tested region.

### Transactivation assay

*TCP4*/*LANCEOLATE* coding region and the genomic fragment of 760 bp of *OVATE*, which includes part of the second intron and second exon (in which the *TCP4/LANCEOLATE* binding site is located), were amplified using the oligos listed in the Table S1. *TCP4*/*LANCEOLATE* coding region was used to generate the *p35S::LANCEOLATE* effector construct. *OVATE* genomic fragment was digested with *NcoI* and *BamHI* restriction enzymes and cloned into the multicloning site of pGreenII 0800 LUC (Hellens et al., 2005) to generate the reporter construct. Leaves of 6-week-old *N. benthamiana* plants were coinfiltrated with a combination of both effector and reporter constructs in equal volumes. Three days after coinfiltrating, Firefly and Renilla Luciferase activities were measured using Dual-Luciferase Reporter Assay System (Promega), following the manufacturer’s instructions. Absolute relative light units (RLU) were measured by GloMAX discover system (Promega). Data were analysed as Luciferase/Renilla ratio and compared with the ratio of the leaves infiltrated with *A. tumefaciens* harbouring the pK7WG2.0 empty vector (control). Four biological replicates and two technical replicates were analysed.

### Ovary and fruit shape analysis

Fruit or ovary shape index is the ratio of the maximum height length to maximum width of a fruit or ovary. Full size mature fruits or ovaries at anthesis were photographed at 300 dpi and the maximum length and width of ovaries/fruit were measured using ImageJ software (https://imagej.nih.gov/ij/). At least five plants of each genotype were analysed, and the average values each taken from 8 to 10 fruits or ovaries per plant were analysed with Tukey’s HSD test (*P* < 0.05) or Student’s t-test (two-tailed).

## ACKNOWLEDGMENTS

We thank the members of Dr Nogueira’s laboratory for helpful discussions. We also thank Americo J.C. Viana, Michel Vincentz, Marcela M. Notini, Cassia R. Figueiredo, and Raphael D. Fava for excellent technical assistance. The authors declare no competing financial interests.

## FUNDING

A.C.Jr was the recipient of The São Paulo Research Foundation (FAPESP) fellowship (no. 2015/07171-0). This work was supported by FAPESP (grant nos. 15/17892–7 and 18/17441–3), and partially by the Brazilian National Council for Scientific and Technological Development (grant no. 409186/2016-3).

## AUTHOR CONTRIBUTIONS

A.C.Jr and F.T.S.N. were responsible for the conception, planning and organization of the experimental time line. A.C.Jr, L.F.F., M.H.V., E.M.S., M.L, and C. S. carried out experiments. A.C.Jr, L.F.F., M.H.V., and F.T.S.N. supervised the development of the experimental plan. F.T.S.N. and R.S. directly supervised the experimental work done by A.C.Jr, L.F.F., and M.H.V. L.E.P.P. and N.O. provided genetic material and helped to analyze data. A.C.Jr, L.F.F., M.H.V., and F.T.S.N. discussed the resulting data. The manuscript was written by A.C.Jr and F.T.S.N.

## Supplemental data

**Fig. S1.** *TCP4/LANCEOLATE* de-repression leads to modifications in ovary and fruit morphology.

**Fig. S2.** MiR319-targeted *TCP4/LANCEOLATE* controls fruit shape in the hybrid backgrounds.

**Fig. S3.** Opposing expression profiles of *TCP4/LA* and *OVATE* during young flower development.

**Fig. S4.** The loss of tomato *CINCINNATA-like TCP* function leads to sepal overgrowth.

**Fig. S5.** Abaxial ectopic expression of a miR319-resistant allele of *TCP4/LA* (*La-2*).

**Fig.S6.** *OVATE* modulates fruit morphology dependent and independently of the miR319/*LANCEOLATE* regulatory hub.

**Fig. S7.** *TCP4/LANCEOLATE* de-repression leads to increase pedicel growth.

**Fig. S8.** *AtPIN1-GFP* expression in tomato gynoecium.

**Fig.S9.** Auxin-associated genes are misexpressed in flowers buds of *ovate* mutant.

**Fig.S10.** *Arabidopsis* miR319-targeted *CIN-TCPs* modulate auxin responses in developing gynoecia.

**Table S1.** Putative TCP4/LANCEOLATE-binding motifs

**Table S2**. Oligonucleotide sequences used in this work.

## Notes

### Competing Interest Statement

The authors have declared no competing interest.

