## Supplemental Figures and Table 1, 2 for "miR319-targeted *TCP4/LANCEOLATE* directly regulates *OVATE* and auxin responses to modulate tomato gynoecium patterning and fruit morphology"

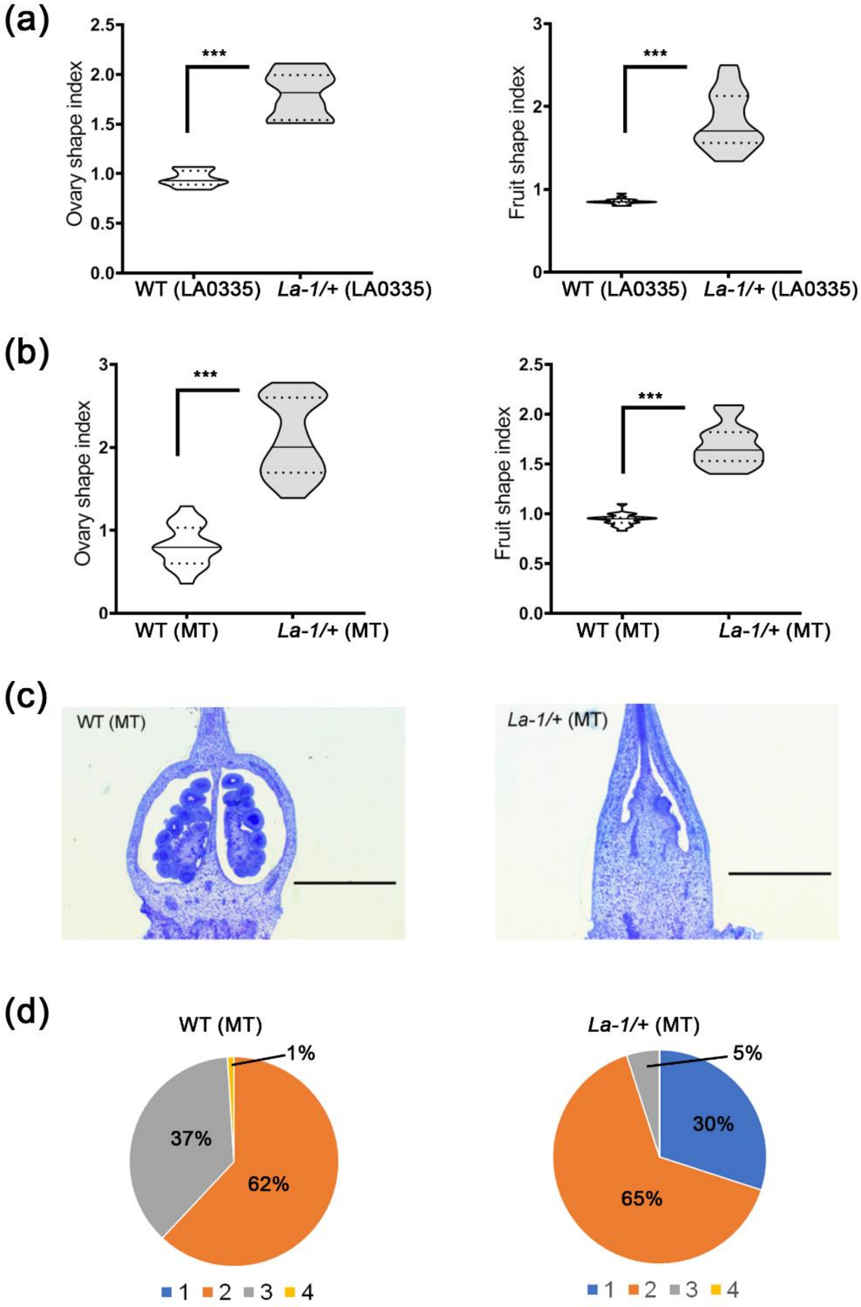


**Fig. S1. *TCP4/LANCEOLATE* de-repression leads to modifications in ovary and fruit morphology.** Violin plots of fruit or ovary shape indexes in LA0335 (**a**) and MT (**b**) backgrounds. The black line indicates the median and the dashed lines indicate the quartiles. (***$P$<0.001). **c**, Longitudinal sections of WT (MT) and *La-1/+* (MT) pre-anthesis ovaries. Bars = 1 mm. **d**, Locule number (1, 2, 3 or 4) percentage in WT (MT) and *La-1/+* (MT).

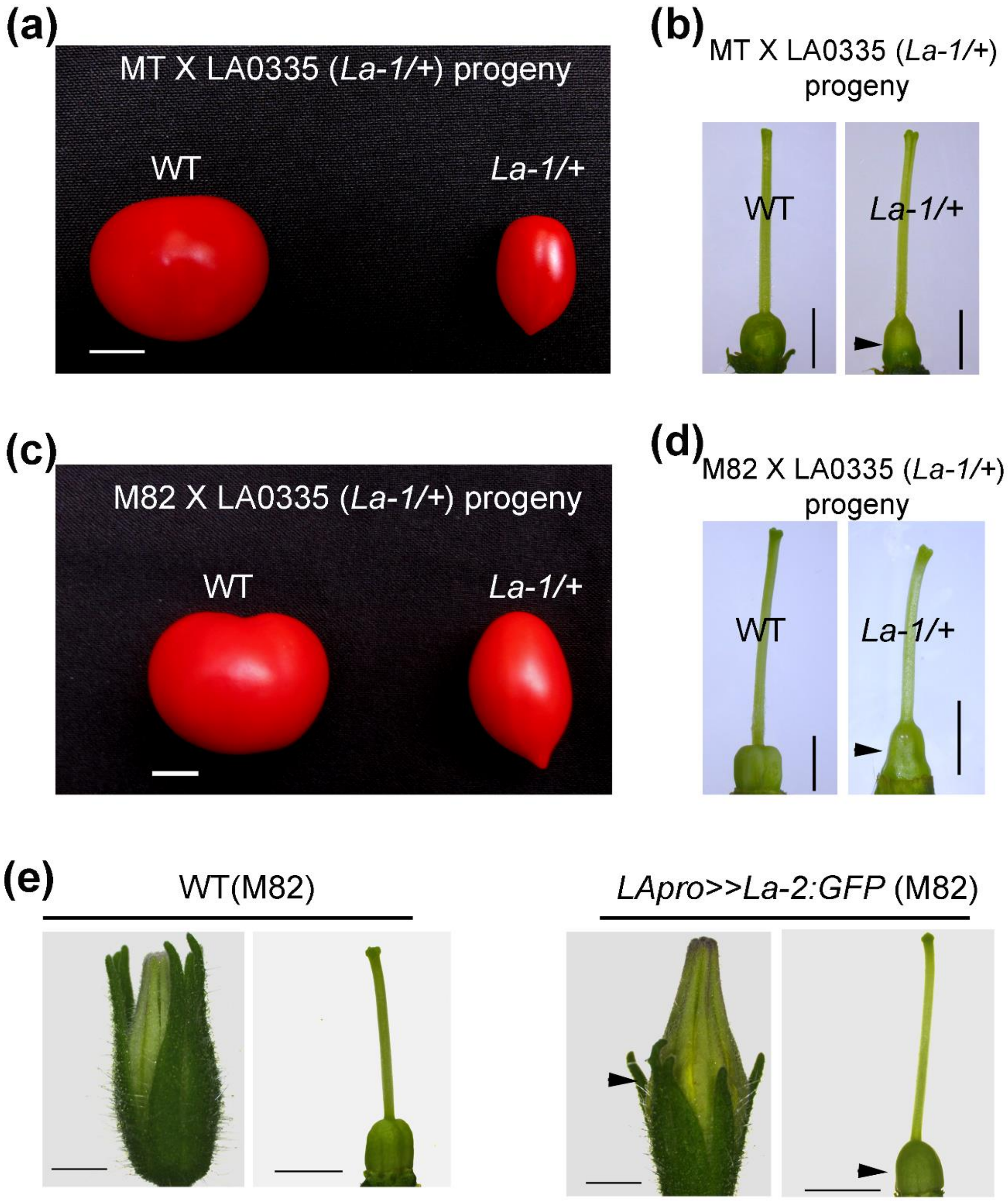


**Fig. S2. MiR319-targeted *TCP4*/*LANCEOLATE* controls fruit shape in the hybrid backgrounds.** In **a-d**, Representative F1 offspring gynoecia and fruits from the crossing between the semi-dominant *Lanceolate* (*La-1/+*; accession LA0335) and 'Micro-Tom' (MT) and M82 backgrounds. Bars = 1 cm (**a, c**) and 2 mm (**b, d**). **e**, Representative flowers and gynoecia of WT (M82) and *LApro>>La-2:GFP* (M82) plants. Arrowheads indicate small sepals in flower and medial constriction in the ovaries. Bars = 2 mm.

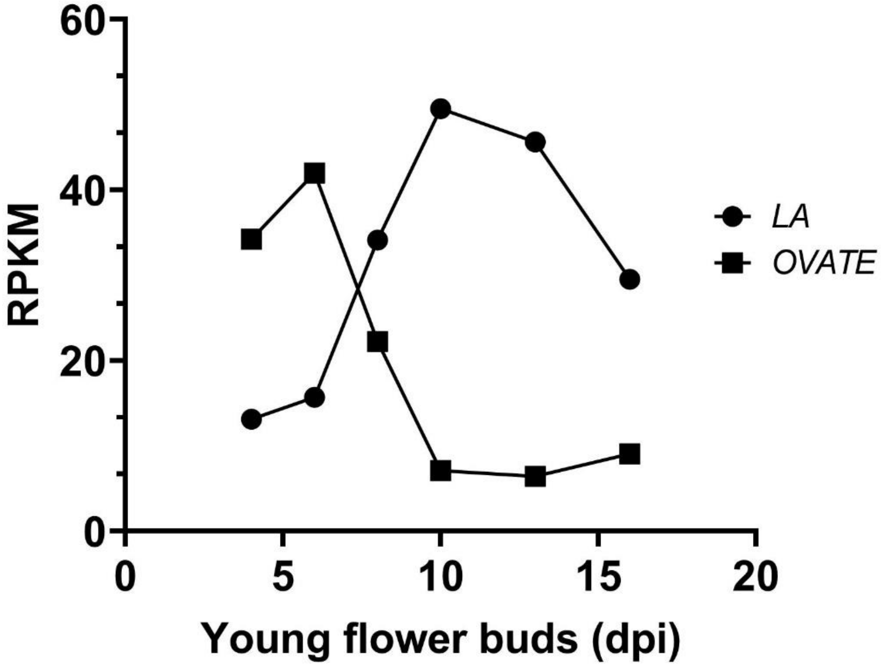


**Fig. S3. Opposing expression profiles of *TCP4/LA* and *OVATE* during young flower development**. Expression patterns of *TCP4/LANCEOLATE* (*LA*) and *OVATE* in RPKM (Reads Per Kilobase Million) values. The data were retrieved from the Transcriptome profiling of young flower buds of WT, *sun, ovate and fs8.1* NILs in the LA1589 background (http://ted.bti.cornell.edu/cgi-bin/TFGD/digital/experiment.cgi?ID=D005).

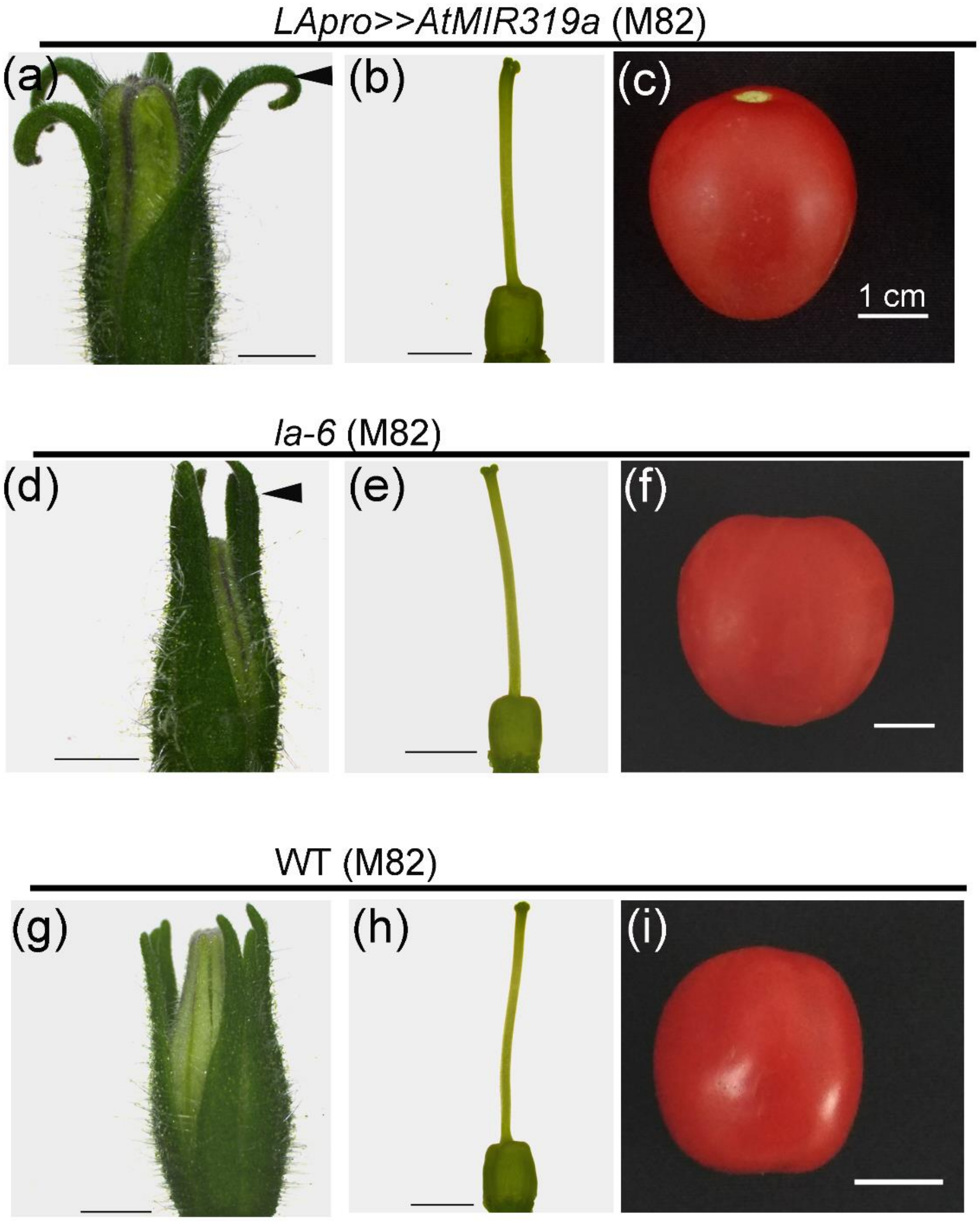


**Fig. S4. The loss of tomato *CINCINNATA-like TCP* function leads to sepal overgrowth.** (a-c**)** Representative imagens of *LApro>>AtMIR319a* (M82) pre-anthesis flower (a), gynoecium (b), and fruit (c). (d-f) Representative imagens of *la-6* (M82) pre-anthesis flower (d), gynoecium (e), and fruit (f). (g-i) Representative imagens of WT (M82) pre-anthesis flower (g), gynoecium (h), and fruit (i). Arrows in **a, d** indicate sepal overgrowth. Bars = 2 mm (**a, b, d, e, g, h**) and 1 cm (**c, f, i**).

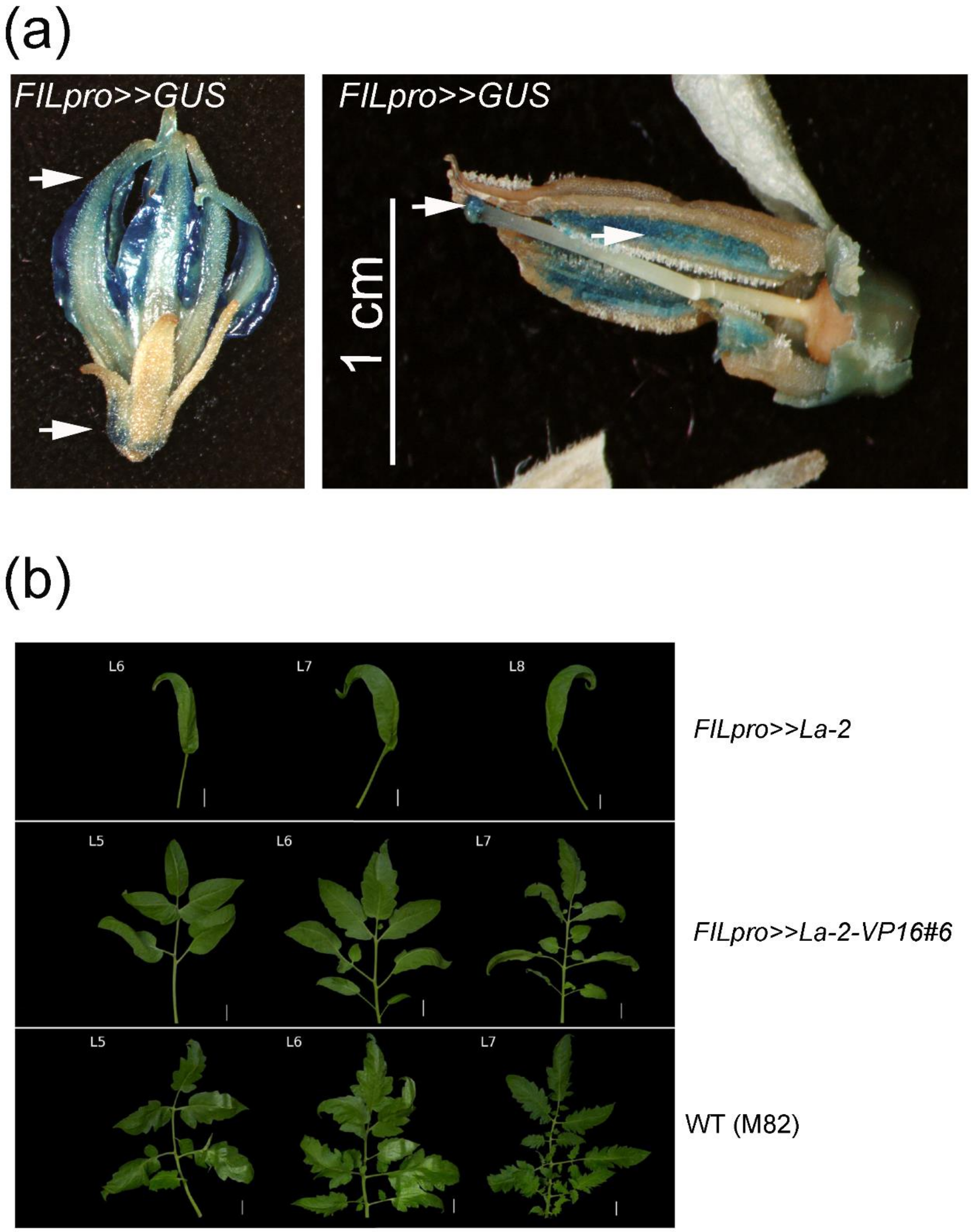


**Fig. S5. Abaxial ectopic expression of a miR319-resistant allele of *TCP4/LA* (*La-2*).** (a) *GUS* expression transactivated by the *FILAMENTOUS FLOWER* (*FIL*) promoter in tomato flower. Arrows indicate abaxial *GUS* expression. (b) Representative leaf (L) imagens of *FILpro>>La-2* (Ori et al., 2007), *FILpro>>La-2-VP16#6*, and WT (M82). Bars = 1 cm.

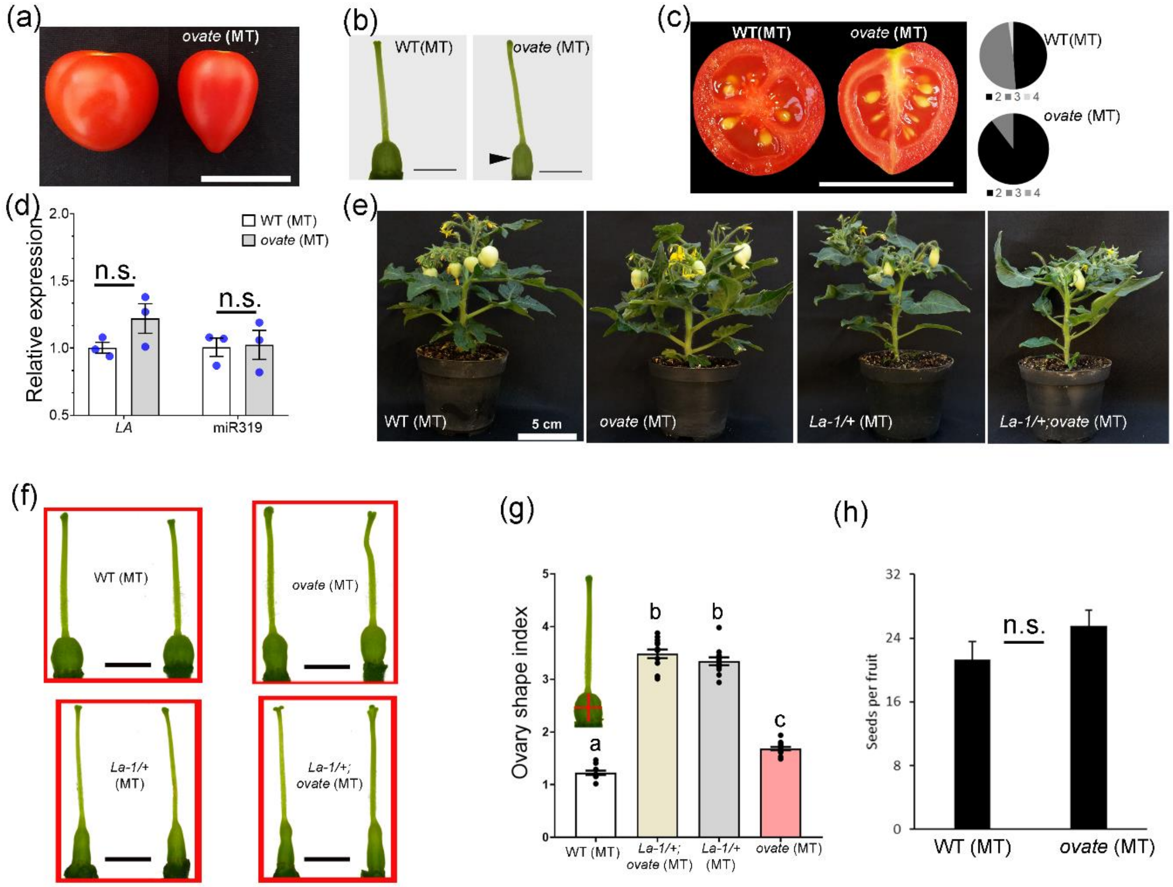


**Fig. S6. *OVATE* modulates fruit morphology dependent and independently of the miR319/*LANCEOLATE* regulatory hub.** (a) Representative imagens of WT (MT) and *ovate* (MT) fruits. Bar = 1 cm. (b) Representative imagens of WT (MT) and *ovate* (MT) gynoecia at anthesis. Arrowhead indicates slight constriction in the ovaries. Bars = 2 mm. (c) Right panel: opened fruits of WT (MT) and *ovate* (MT) showing placenta and seeds. Bar = 2 cm. Left panel: locule number (2, 3, or 4) of WT (MT) and *ovate* (MT) fruits. (d) Relative expression levels of *TCP4*/*LANCEOLATE* (*LA*) and miR319 in 5-8 days post-inflorescence (dpi) floral buds (n =3). **P*<0.05 according to Student's t-test (two-tailed). Values are means ± s.e. (e) Representative imagens of 52-days-old plants. (f) Representative gynoecia at anthesis of WT (MT), *ovate* (MT), *La-1/+* (MT), and double mutant *La-1/+;ovate* (MT). Scale bars = 2 mm. (f) Ovary shape index is the ratio of length to width (red cross). Values are mean ± s.e. Letters indicate the significant differences among different genotypes evaluated by Tukey's HSD test (*P*< 0.05).

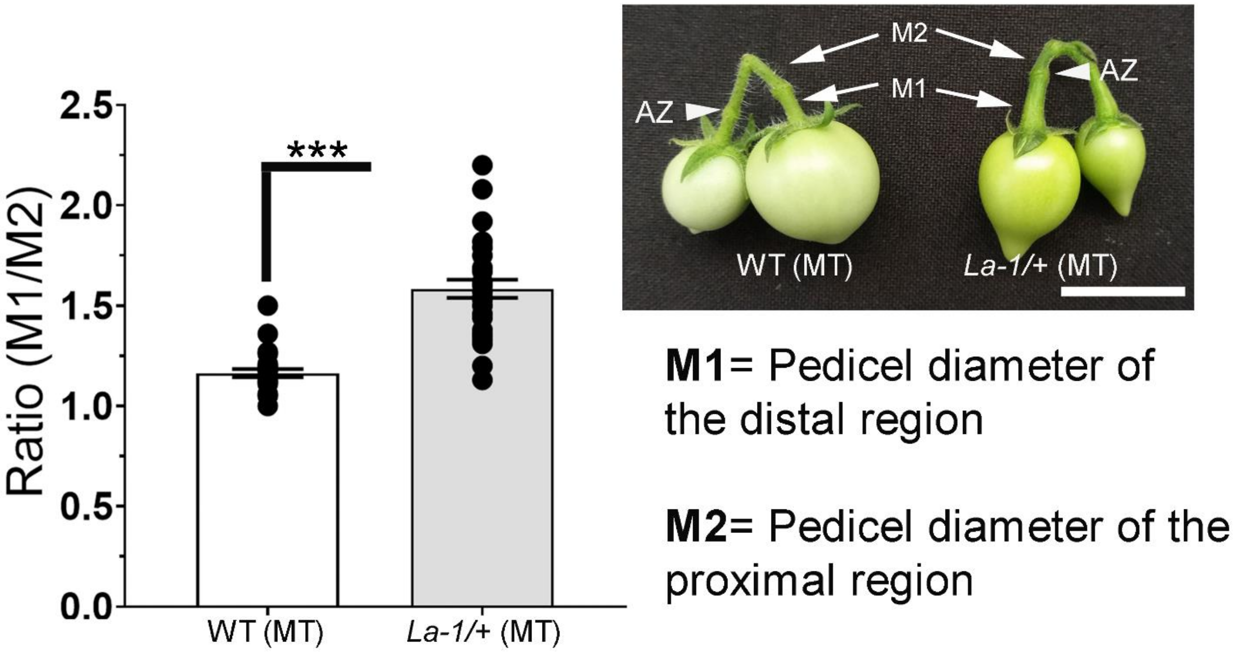


**Fig. S7.** ***TCP4/LANCEOLATE*** **de-repression leads to increase pedicel growth.** Ratio of pedicel growth (n =15). ****P*<0.001 according to Student's t-test (two-tailed). Values are means ± s.e. AZ, abscission zone. Bar = 2 cm.

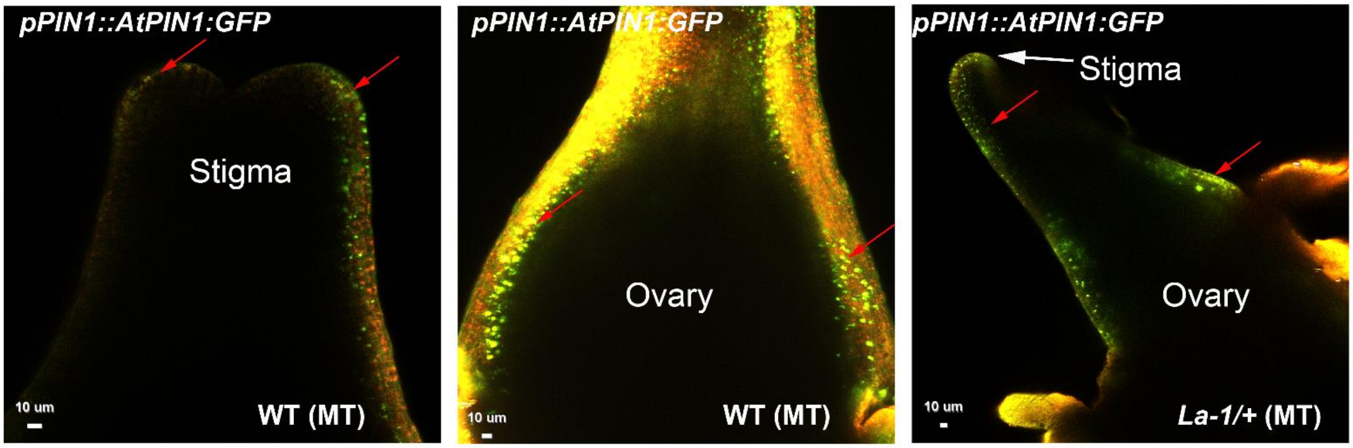


**Fig. S8. *AtPIN1-GFP* expression in tomato gynoecium.** Confocal merged images were obtained by merging a GFP-filtered image with an identical RFP-filtered image. Arrows indicate polarized AtPIN1-GFP towards the tip and AtPIN1-GFP proteins localized in brefeldin A (BFA) body-like structures. Bars = 10 μm.

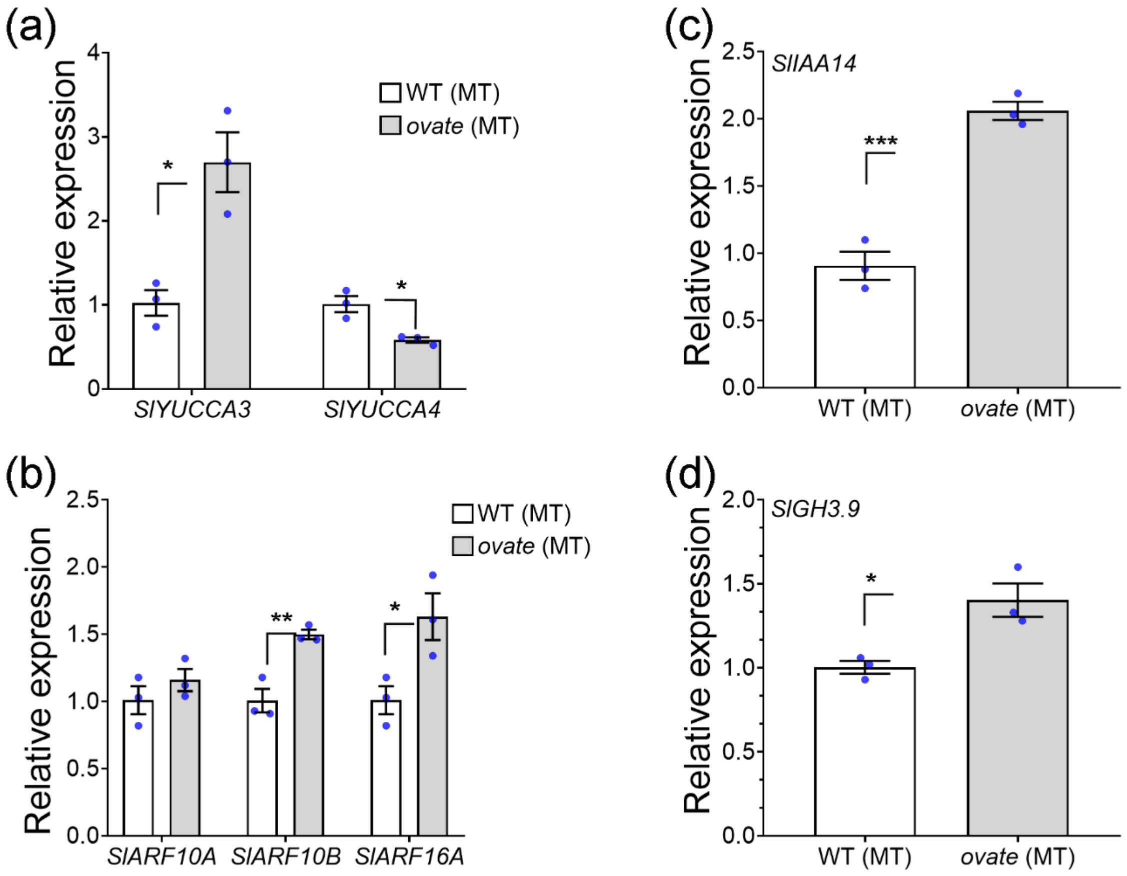


**Fig S9. Auxin-associated genes are misexpressed in flowers buds of *ovate* mutant.** Relative expression levels of *SlYUCCA3/4* (a), *SlARF10A/B* and *SlARF16A* (b), *SlIAA14* (c), and *SlGH3.9* (d) in 5-8 days post-inflorescence (dpi) floral buds of WT (MT) and *ovate* (MT) (n =3). **P*<0.05, ***P*<0.01, ****P*<0.001 according to Student's t-test (two-tailed). Values are means ± s.e.

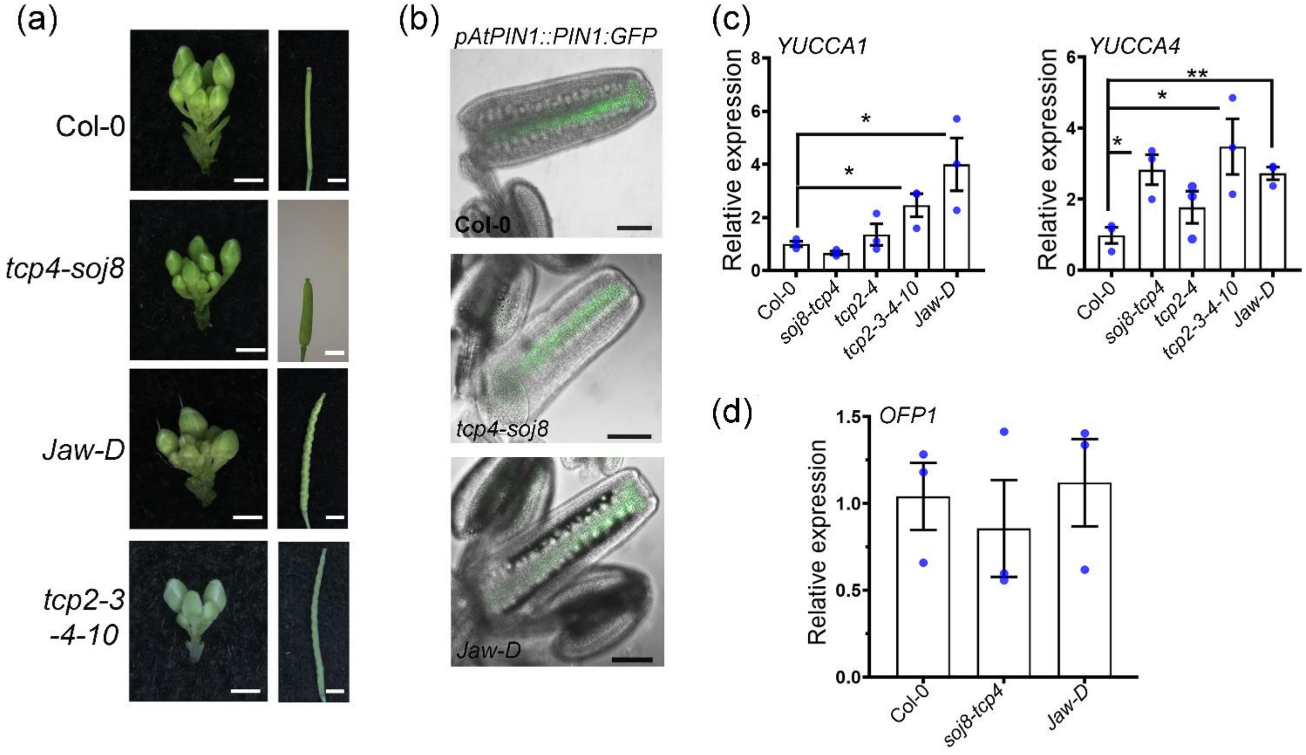


**Fig. S10.** ***Arabidopsis*** **miR319-targeted** ***CIN-TCPs*** **modulate auxin responses in developing gynoecia.** (a) Representative imagens of inflorescence and fruits of *Arabidopsis* genotypes. Bars = 1 mm (inflorescences) and 2 mm (siliques). (b) Representative imagens of developing gynoecia from *Col-0; pAtPIN1::PIN1-GFP*, *tcp4-soj8; pAtPIN1::PIN1-GFP*, and *Jaw-D; pAtPIN1::PIN1-GFP*. Bars = 100 µm. Relative expression levels of *YUCCA1* (*YUC1*) and *YUC4* (c) and *OVATE FAMILY PROTEIN1* (*OFP1*) (d) in floral buds at 1-7 developmental stage (n =3). **P*<0.05, ***P*<0.01 according to Student's t-test (two-tailed). Values are means ± s.e.

**Supplementary Table S1. Putative TCP4/LANCEOLATE-binding motifs**

| Gene | ID | Motif | Strand | Position | *Score |
|---|---|---|---|---|---|
| ***OVATE*** | Solyc02g085500 | T[GGACC]AA | + | -4078 | 0.93 |
| | | A[GGACC]CG | + | +1372 (ORF) | n.d. |
| ***SlARF10B*** | Solyc06g075150 | A[GGACC]AT | + | -1265 | 0.93 |
| ***SlYUCCA4*** | Solyc06g008050 | G[GGACC]TC | + | -893 | 0.87 |

*Score of putative TCP4-like binding sites from The Plant Promoter Analysis Navigator (PlantPAN 3.0; http://PlantPAN.itps.ncku.edu.tw). n.d., no determined.

**Supplementary Table S2.** Oligonucleotide sequences used in this work.

| Name | Sequence (5'-3') | Purpose | * Accession number | Reference |
|---|---|---|---|---|
| Hyb-LA - F | CCAGCCCACTTTGGAGGAAA | ISH (RNA probe) | Solyc07g062680 | Ori et al., 2007 |
| Hyb-LA - R | TTCAAGTAGAGCATTTCCCTGTCA | | | |
| miR319a | AACCTGACTTCCCTCGAGGGA | ISH (LNA probe) | MI0009978 | this work |
| scrambled miRNA | GTGTAACACGTCTATACGCCCA |  | ------------ |  |
| chipOVATE - F | GGAAAGTGAAGGAGAGTTTCGC | ChiP-qPCR | Solyc02g085500 | this work |
| chipOVATE - R | GCTGTTCCAGCTCATTCTTCTC | | | |
| proOVATE (R1) - F | GTCATACTCAGCAAGTGTCC | ChiP-qPCR | Solyc02g085500 | this work |
| proOVATE (R1) - R | CAGTTGGATTGCATGTGTTAC | | | |
| proSlARF10B -F | ATGGCTCCCATGCCTTGTAC | ChiP-qPCR | Solyc06g075150 | this work |
| proSlARF10B -R | GAACGATAACGAACTCCACAGG | | | |
| proSlYUCCA4 -F | GACATGTGTTTAAGACGTTC | ChiP-qPCR | Solyc06g008050 | This work |
| proSlYUCCA4 -R | CATGAGACATAGTACTCTAAATATG | | | |
| Luc-LA - F | CACCATGGCAGAAACGTCAA | Transactivation assay | Solyc07g062680 | this work |
| Luc-LA - R | TCAATGGCGAGAATCAGAGG | | | |
| lucOVATE - F | CGCGGATCCGCGATCCACATTAAGAGACGCTG | Transactivation assay | Solyc02g085500 | this work |
| lucOVATE - R | CATGCCATGGCATGTCATTTCCATCATCGATCTC | | | |
| qLA-F | TGCAGCAGCTATTCGGTCAA | RT-qPCR | Solyc07g062680 | this work |
| qLA-R | ACCCAGAGAATCCGCCTACT | | | |
| | | | MI0009978 | |
| RT-SlmiR319a | GTCGTATCCAGTGCAGGGTCCGAGGTATTCGCACTGGATACGACGGAGCT | Stem-loop RT-qPCR | | this work |
| miR319 F | GCATAGCTTGGACTGAAGGG | Stem-loop RT-qPCR | MI0009978 | this work |
| Reverse universal | GTGCAGGGTCCGAGG | Stem-loop RT-qPCR | | Varkonyi-Gasic et al., 2007 |
| qOVATE-F | GCTGATGAGTACAGCGGGAT | RT-qPCR | Solyc02g085500 | this work |
| qOVATE-R | CTTGTGACGACGCCTCATTG | | MI0009978 | |
| qSlOFP20-F | AGCAGCAGAAGACATAGCCAC | RT-qPCR | Solyc10g076180 | this work |
| qSlOFP20-R | ATGTGGACGTTGTGATCGAGTT | | | |
| qSlYUCCA1-F | GTACTCGACGTTGGAGCATTATC | RT-qPCR | Solyc06g065630 | This work |
| qSlYUCCA1-R | TGAAGAAATCATTTCCCTTAAACC | | | |
| qSlYUCCA3-F | TGGAGAAATACAAGAAATTGATTG | RT-qPCR | Solyc09g091090 | This work |
| qSlYUCCA3-R | GTACTCGACGTTGGAGCATTATC | | | |
| qSlYUCCA4-F | TCGATTCTGTTCTTCTTGCTACT | RT-qPCR | Solyc06g008050 | This work |
| qSlYUCCA4-R | CTGATAGTCCTCTTCTTGTGAAA | | | |
| qSlYUCCA5-F | GGCACCGTTGAACTTGTCAC | RT-qPCR | Solyc06g083700 | This work |

| | | | | |
|---|---|---|---|---|
| qSlYUCCA5-R | CCTTTCCAATTATTTGGGATTT | | | |
| qSlYUCCA6-F | CCAGAAGAAGGACCATTTGC | RT-qPCR | Solyc09g074430 | This work |
| qSlYUCCA6-R | TACCATTTTCCACCACAACATC | | | |
| qSlARF10A-F | TCACACAACTCACCACCAAG | RT-qPCR | Solyc06g075150 | This work |
| qSlARF10A-R | AGGCGGATAACGAGAATAAC | | | |
| qSlARF17-F | TTATCTGTTCGTTTTCTCGC | RT-qPCR | Solyc11g013480 | This work |
| qSlARF17-R | TCCCTCATAGTAACTTCATC | | | |
| qSlIAA14-F | GGGTTTTCTGAGACTGTTG | RT-qPCR | Solyc09g083290 | This work |
| qSlIAA14-R | AGGTGGCTTGATTGGATC | | | |
| qSlGH3.4-F | CGTGATGAATCTGTATGTGCCTGG | RT-qPCR | Solyc02g092820 | This work |
| qSlGH3.4-R | GTTAGGACTGGACGTGCAACAAG | | | |
| qSlGH3.9-F | CTCTGCTCTCAGCCCATCTCTG | RT-qPCR | Solyc07g063850 | This work |
| qSlGH3.9-R | GCTGTGATCTTCTCGCAAGTTCC | | | |
| OVATE_GenF<br>OVATE_GenR | GCGTGTGGAATTTGGAGAGGACAGA<br>GTGGAGTTAGAAGTCCCATGAGCAAG | Genotyping | Solyc02g085500 | this work |
| qAtYUCCA1-F<br>qAtYUCCA1-R | TGGGATGCCGAAAACGCCGT<br>CGTCAGACGCCGTTCCAAGGA | RT-qPCR | At4g32540<br>MI0009978 | Lucero et al., 2015 |
| qAtYUCC4-F<br>qAtYUCC4-R | CTTTCGGGAACAGCCTATGAC<br>GGATTTATTGAAATGAAGATG | RT-qPCR | At5g11320 | this work |
| qAtOFP1-F<br>qAtOFP1-R | CAAGTTATCCAAAACCGCAACCT<br>CACCGAGATCCGCGGCGACCG | RT-qPCR | At5g01840 | this work |
| qAtPP2A-F<br>qAtPP2A-R | CCTGCGGTAATAACTGCATCT<br>CTTCACTTAGCTCCACCAAGCA | RT-qPCR | At4g25420<br>At1g13320<br>At1g13320 | this work |

*: All accession numbers can be found in the Sol Genomics Network, miRbase (22 release) or the or the National Center for Biotechnology Information (NCBI).
